# Rare Monoclonal Antibody Discovery Based on Indirect Competitive Screening of Single Hapten-specific Rabbit Antibody Secreting Cell

**DOI:** 10.1101/2022.04.14.488422

**Authors:** Yuan Li, Peipei Li, Yuebin Ke, Xuezhi Yu, Wenbo Yu, Kai Wen, Jianzhong Shen, Zhanhui Wang

## Abstract

Rare antibody that is able to tolerate physio-chemical factors is preferred and highly demanded in diagnosis and therapy. Rabbit monoclonal antibodies (RmAbs) are distinguished owing to their high affinity and stability. However, the efficiency and availability of traditional methods for RmAb discovery are limited, especially for small molecules. Here, we present an indirect competitive screening method in nanowells, named CSMN, for single rabbit antibody secreting cells (ASCs) selection with 20.6 h and proposed an efficient platform for RmAb production against small molecule with 5.8 days for the first time. Chloramphenicol (CAP) as an antibacterial agent has the great threats for public health. We applied the CSMN to select CAP-specific ASCs and produced one high affinity RmAb, surprisingly showing extremely halophilic properties with an IC_50_ of 0.08 ng mL^-1^ in saturated salt solution which has as yet not been shown by other antibodies. Molecular dynamic simulation showed that the negatively charged surface improved the stability of the RmAb structure with additional disulfide bonds compared with mouse antibody. Moreover, the reduced solvent accessible surface area of the binding pocket increased the interactions of RmAb with CAP in a saturated salt solution. Furthermore, the RmAb was used to develop an immunoassay for the detection of CAP in real biological samples with simple pretreatment, shorter assay time, and higher sensitivity. The results demonstrated that the practical and efficient CSMN is suitable for rare RmAb discovery against small molecules.

## Introduction

Monoclonal antibodies (mAbs) are essential tools in therapeutic, diagnostic, and biological filed, predominantly based on the mouse monoclonal antibodies (MmAbs). Nowadays, there has been a tendency of using rabbit monoclonal antibodies (RmAb) owing to their high affinity, specificity, and stability ^1^. Hapten-specific mAbs are important tools in drug detection, food analysis, and environmental monitoring. Rabbit produces more diverse and stable antibodies against more antigens, including small molecules, due to their specific immune mechanism and unique antibody structure ^2^. Thus, RmAb provide the dawn to meet the requirement of outstanding reagents for laboratory research and increasingly need for diagnostic and therapeutic applications.

RmAb was firstly produced by hybridoma technology as MmAbs since 1995^3^, however, developing rabbit hybridomas was rather complicated and lower efficient owing to the unstable rabbit myelomas compared with classic mouse myelomas. Therefore, efforts were originally made to generate rabbit-mouse hybridomas which was also not so successful due to the poor fusion efficiency, genetic instability and decrease in antibody secretion^4^. Immortalization of peripheral B cells with Epstein-Barr virus is another method to induce long-term growth of B cells for antibody production^5^, but the low efficiency of this method has seriously obstructed the efficiency of RmAb preparation. To overcome these issues, several display platforms have been introduced to discovery RmAb combing with the recombinant expression technology, such as yeast surface display, ribosome display, and phage display^6, 7^. However, the variable domain genes of an antibody are randomly combined during the construction of antibody libraries, leading to the natural cognate pairing of heavy chain and light chain lost. The mis-paring of the heavy chain and light chain is the main bottleneck to antibody discovery, especially for small molecules^8^. Thus, exploring of RmAb discovery methods is still on the road and urgently needed.

With advances of fluorescence-activated cell sorting (FACS) and microfluidic techniques, some studies have focused on screening of antigen-specific B cells from a large population of primary B cells and produced single cell antibody with natural light- and heavy-chain pairs. Chen et al. and Kramer et al. have produced RmAb recognizing SARS-CoV-2^9^ and HLA-DR ^10^, respectively, from single memory B cells screened by FACS based on the cell surface IgG. Antibodies are directly generated by the antibody-secreting cells (ASCs) during the immune response. Thus, comprehensive evaluation and precise selection of antigen-specific ASCs would be beneficial to antibodies discovery. Identifying antigen-specific ASCs by cell surface staining is not feasible owing to the lack of IgG on the ASC surface. To resolve this issue, Ramirez et al. demonstrated heterofunctional particles as single cell sensors to capture secreted antibodies and isolate antigen-specific ASC by FACS ^11^. However, this approach is not suitable to select ASCs from rabbit, due to the ASC surface markers of rabbit are less known. Droplet-based microfluidic technologies are introduced to encapsulate single ASC in a given volume to identify the secreted mAbs ^12^. However, the frequent false-negative rate and limited microdroplet volume significantly interfered the screening efficiency of single ACS.

Recently, it has been demonstrated that secreted proteins from single cells can be analyzed with fluorescent-based assays in nanowells or microwells, which can be applied for the screening of hybridomas ^13^ and human ASCs ^14^. Various direct staining methods have been developed to identify the ASCs that secrete antibodies of interest. Love et al. applied the ovalbumin-coated slides in microwells to capture the secreted anti-ovalbumin antibody from the corresponding hybridoma and realized an effective screening of hybridomas ^15^. Unlike proteins, small molecules that cannot be directly coated in the microwell and always chemically conjugated to proteins as the immunogens or coating antigens due to the small size. With these chemical modifications, not only target molecules alone but also linkers of haptens and amino acid residues on the protein are potential epitopes that can induce the generation of ASCs. Therefore, the strategies that could precise select the ASCs specific to proteins are not yet available to small molecules.

In this study, we proposed a competitive screening method, named CSMN, for hapten-specific rabbit ASC selection based on nanowells. Chloramphenicol (CAP) is used as a model analyte, which is widely used for the prevention and treatment of poultry and aquatic diseases. Due to the animal-derived foods with CAP residue posing great health hazards to humans ^16, 17^, the use of CAP has been totally banned worldwide. Antibody-based analytical methods, i.e., the immunoassays, usually are thought to be preponderant, with the characteristics of high sensitivity, reproducibility, short analysis time, and simple sample pretreatment^18^. However, these advantages do not always fully realize their potential, and suffer from decreased antibody activity in harsh detection environments containing high concentrations of salt, solvent, acid, and alkali substances. Recently, an indirect competitive enzyme-linked immunosorbent assay (icELISA) based on MmAb was developed for the CAP detection, which lacks sufficient affinity to meet the requirement of trace CAP detection with multiplex sample pretreatment ^19^. Thus, RmAb with high affinity, stability, and specificity is obviously beneficial for detecting CAP at low concentrations in complicated biological samples.

## Results and discussions

### Procedure of the CSMN and production of single cell RmAb

Fig. 1 shows a detailed flowchart of the CSMN and single cell RmAb production in this study. The nanowell surface was first modified by coating with CAP-BSA for 12 h, and then the splenocytes from the immunized rabbit mixed with PE-anti-rabbit F(ab’)_2_ antibody, and were evenly distributed into the nanowells at single cell level with 10 min. Next, the RmAb secreted by single ASCs in the individual nanowells for 8 h could be captured by the coated CAP-BSA and stained with PE-anti-rabbit F(ab’)_2_ antibody. At 0-4 h, the high fluorescence intensity of the nanowell indicated the secreted antibody from the loaded ASC could recognize the CAP-BSA. Other nanowells loaded with non-CAP-BSA-specific ASC and non-ASC showed no detectable fluorescence. Next, CAP was added to the nanowells as a competitor with CAP-BSA for the identification of the CAP-specific ASC at h 4 for a further incubation. The ASCs loaded in the nanowells with significantly decreased fluorescence intensity at 4-8 h were selected. Other nanowells with a continuous increase in fluorescence intensity due to the secreted antibodies of these loaded ASCs were BSA-specific and these cells were abandoned, which was the key step for the precise identification of ASC against small molecule in the study.

**Fig. 1.**
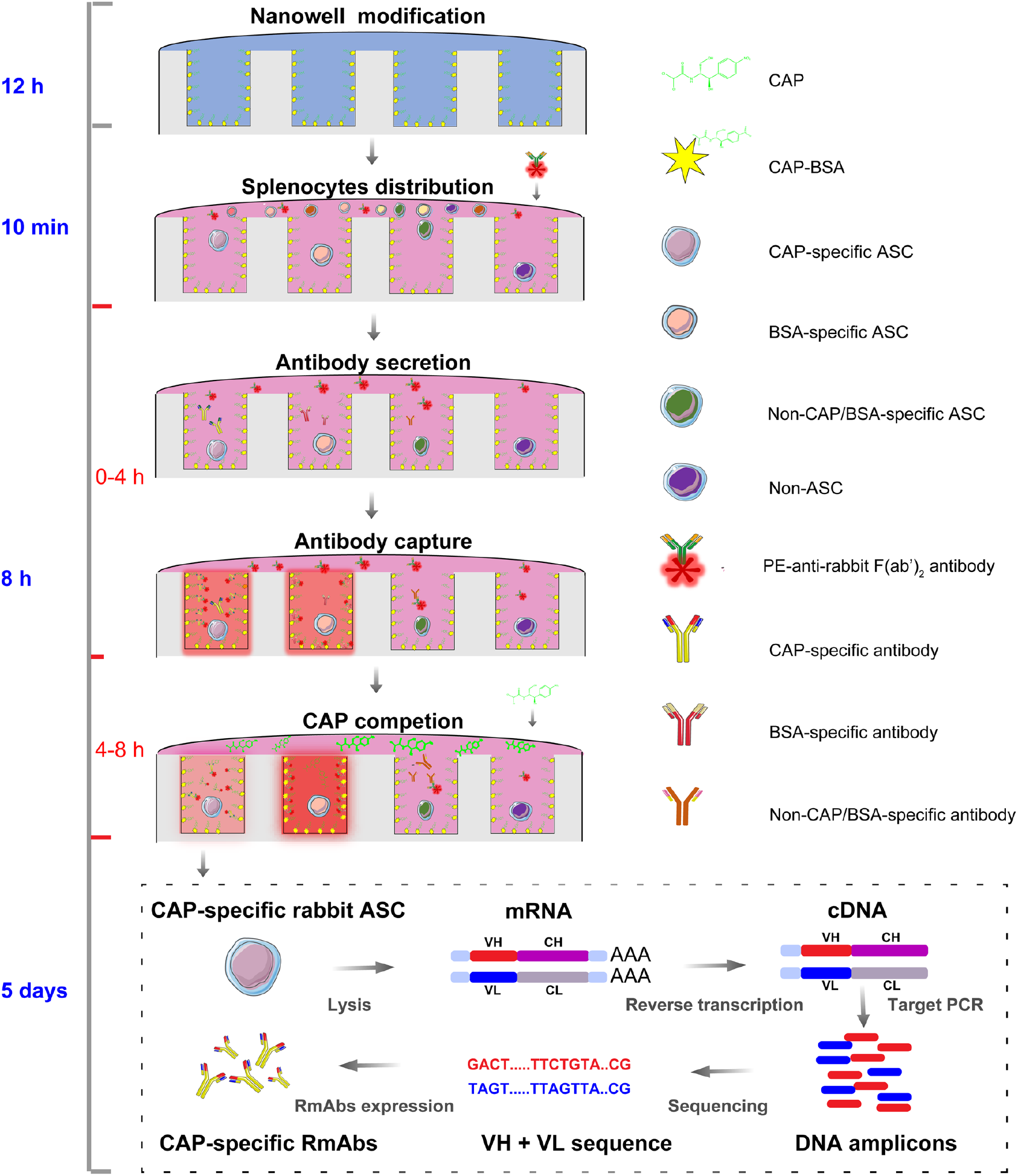
Schematic overview of the competitive screening of single rabbit ASCs by CSMN and the discovery of RmAbs to CAP.

The CAP-specific ASCs were selected and lysed at the single cell level to prepare the antibody mRNA. Then, the mRNA was reverse transcribed to cDNA, and the VH and VL were amplified from cDNA with the specific primers. The DNA amplicons of VH and VL were sequenced by the Sanger sequencing method. CAP-specific RmAbs were expressed *in vitro* by the eukaryotic expression system.

### Optimization of nanowell size and cell numbers in the CSMN

The larger nanowell always contributed to higher cellular occupancy ^20^; however, the single-cell occupancy of the larger nanowell with 37% was significantly lower than that of similar nanowell with 52-92% ^21-23^. Whereas, the retrieve of single cell is severely difficult due to the very small liquid volume when the size of nanowell is similar to the size of the cells ^24^. To ensure both high single-cell occupancy and high retrieve rate, a suitable size of nanowell is critical to for the efficient and precise selection of antigen-specific ASCs. The diameter of rabbit splenocytes has been reported to be approximately 8–13 µm ^25^; thus, nanowells with diameters ranging from 20 µm to 100 µm were firstly optimized in this study.

To test the single-cell occupancy of the nanowell, the number of loaded cells was adjusted to the half of the well numbers (Fig. S1), i.e., 3 × 10^4^ cells mL^-1^ for 100-µm-diameter H100, 2.5 × 10^4^ cells mL^-1^ for 40-µm-diameter U40, 1 × 10^4^ cells mL^-1^ for 25-µm-diameter U25, and 1.5 × 10^5^ cells mL^-1^ for 20-µm-diameter 370K. The Fig. 2A showed that the cellular occupancies of U40 (90.9%) and H100 (72.8%) were higher than that of 370K (67.6%) and U25 with (77.9%). However, the single-cell occupancy of U25 with 54.3% was highest compared to those of other three nanowells all below 46.3% (Fig. 2A). Besides, the UFO design with two auricles of U25 with a 25-µm-diameter provides a high single-cell retrieve efficiency with a volume of 12 pL. Thus, U25 was chosen for the single rabbit ASC selection. The procedure of splenocytes distribution including cell loading, centrifugation and cell plating was showed in Fig. 2B. The number of the loaded rabbit splenocytes in U25 was then optimized with 0.5 × 10^4^ cells (Fig. 2C), 1 × 10^4^ cells (Fig. 2D), and 2 × 10^4^ cells (Fig. 2E). As observed in Fig. 2D and 2E, a larger number of splenocytes gave better capture efficiency; however, excessive splenocytes resulted in multiple-cell occupancy in the nanowells (Fig. 2E). Thus, splenocytes plated on the U25 were at the optimum number of 1 × 10^4^ cells, followed by a brief centrifugation to ensure the cells were loaded in the nanowells as shown in Fig. 2D.

**Fig. 2.**
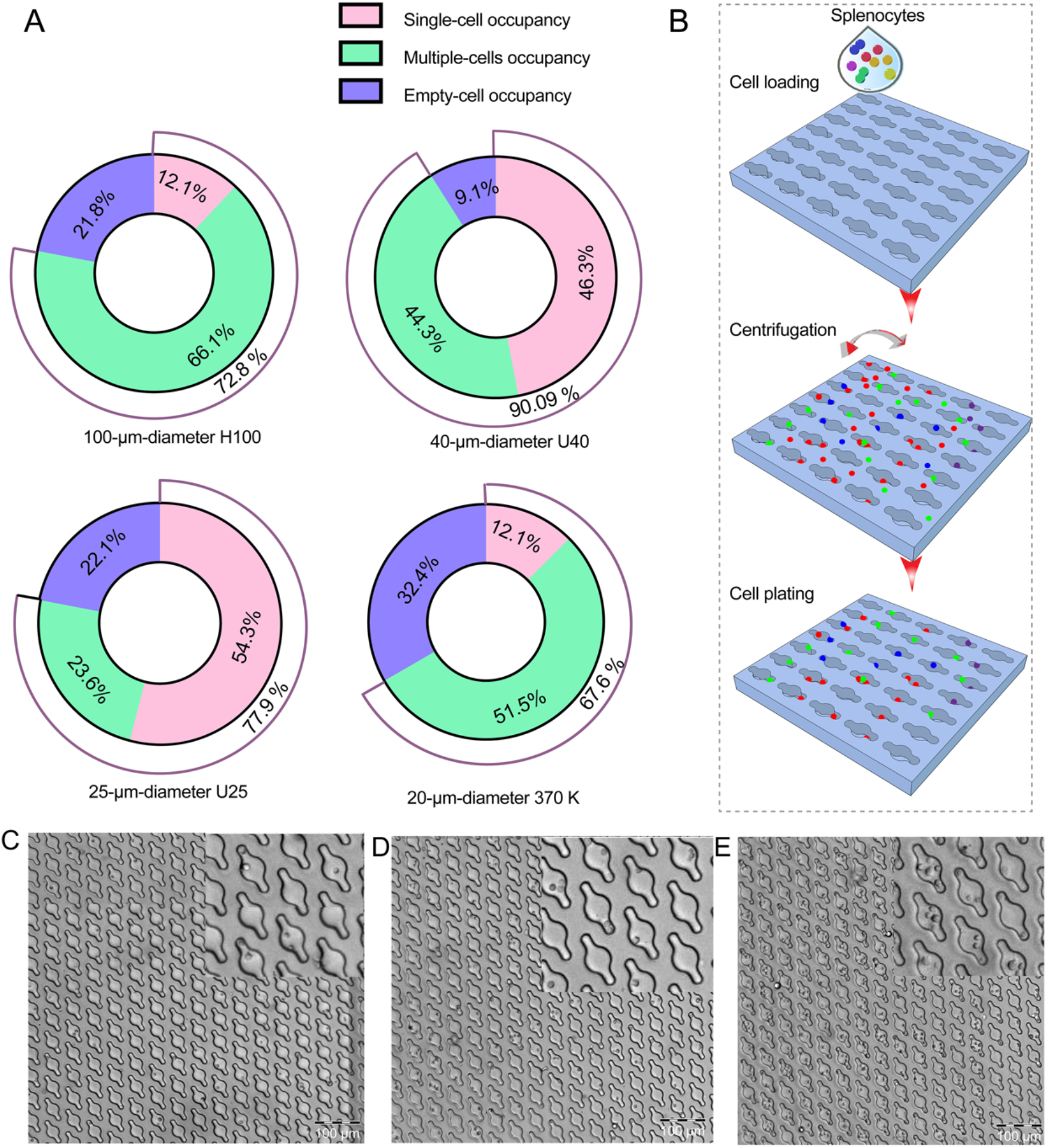
Optimization of the nanowell size and cell numbers. (A) The single-cell occupancy, multiple-cells occupancy and empty-cell occupancy of four nanowells including H100, U40, U25, and 370K. (B) Schematic overview of the splenocyte distribution in U25 with subsequent steps of cell loading, centrifugation, and cell plating. (C) Single-cell occupancy with 0.5 × 10^4^ cells in U25. (D) Single-cell occupancy with 1.0 × 10^4^ cells in U25. E. Single-cell occupancy with 2 × 10^4^ cells in U25.

### Selection of single CAP-specific ASC by the CSMN

Identification and isolation of antigen-specific B cells of interest are crucial steps to discover desired antibody antibodies using single cell techniques. In the FACS, a flow of passing cells is interrogated and antigen-specific B cells of interest are identified and subsequently isolated based on the fluorescently stained cell surface markers. The FACS enables high-throughput isolation of antigen-specific B cells, but only allowing the selection of cells with membrane antibodies such as memory B cells. Since the secreted antibodies from the ASCs could exactly reflect the real affinity of the antibody during the immune stimulation, the precise selection of antigen-specific ASCs is beneficial for high-affinity antibody discovery. Besides, single ASCs secrete thousands of antibodies per second per cell ^26^, which is significantly higher than the number of membrane-bound antibodies on one memory B cell surface limited to around 1 × 10^5 27^, which would afford a more sensitive monitoring. Droplet-based microfluidics capsulizing cells into aqueous droplets holds great potential for functional screening of ASCs at the single-cell level. However, the low single-cell encapsulation efficiencies and physical damage to cells have seriously hindered its wide application ^28^. In this study, CellCelector nanowell array, which combines high-content imaging cytometry system and high-precision single-cell-picking robotics ^29^, for hapten-specific rabbit ASC selection. The roofless nanowell structure permits rapid and direct cell retrieval by capillary and the flexible modification endows the monitoring of secreted antibody based on a tailor-made and effective assay. These assays enable a selection of the best performing ASCs and reduce the cost and time required for the subsequent *in vitro* production and characterization of the obtained antibodies.

Several direct one-step assays focused on determining the affinity of the secreted antibodies toward the coated antigens, have been applied to antibody production based on nanowell for hepatitis C virus ^30^ and ovalbumin ^31^. However, these direct assays are not suitable for precise hapten-specific ASC selection owing to the need for at least two steps to completely identify of the secreted hapten-specific antibody, including the first step of identification of the binding to coating antigen and the second step of identification of binding to target analyte. In this study, we proposed a novel competitive assay to screen the hapten-specific rabbit ASCs based on the U25 nanowell, in which the coated hapten-BSA first bind to the secreted antibodies to induce an increase in fluorescence intensity for the first step identification, which is followed by the addition of target analyte to induce a decrease in fluorescence intensity for the second step identification. CAP was used as a model analyte to identify single ASCs and produce CAP-specific RmAbs by the CSMN. Before the rabbit splenocyte identification procedure, the rabbit antisera was monitored after the sixth immunization. Seven days after the sixth immunization, the rabbit #6 with the best antisera IC_50_ value of 0.27 ng mL^-1^ (Table S1) was sacrificed for the splenocytes collection; and then the immunized splenocytes were submitted to the CSMN.

The PE-anti-rabbit F(ab’)_2_ antibody was used to stain the secreted CAP-specific antibodies from the ASCs in the U25, which is represented by the five regions shown in Fig. S2A–S2C. The mean gray value of the fluorescence intensity of the five regions was significantly increased at 0 h-4 h and slightly changed with the addition of CAP at 4 h-8 h, as observed in Fig. S2D. This result may be due to the insignificant frequency (0.013%) of CAP-specific ASCs in the immunized splenocytes. Besides, the slight change in fluorescence intensity with the addition of CAP indicated that the CAP could indeed compete for the secreted CAP-specific antibodies from the coated CAP-BSA. Then the fluorescence intensity of the individual nanowells containing single CAP-specific ASC was analyzed, as shown in Fig. 3A. We observed that the fluorescence intensity of the nanowell containing CAP-specific ASC significantly increased at 0-4 h and decreased at 4-8 h after the addition of CAP; this was in contrast to the continuous increase in fluorescence intensity observed in the nanowell loaded with BSA-specific ASCs (Fig. 3B), and the lack of fluorescent signal of the nanowell containing the non-CAP/BSA-specific ASC or non-ASC (Fig. 3C). Fig. 3D shows that the CAP competes with CAP-BSA for the CAP-specific antibody; this causes the PE-labeled anti-IgG to be removed from the nanowell, resulting in a decrease in fluorescence intensity (Fig. 3E). This is the most important step to ensure the selection of ASCs that specifically recognize CAP. In the case that the antibody secreted from the ASC is BSA-specific, the BSA-specific antibody could not be competed out from the CAP-BSA by CAP as shown in Fig. 3F, which result in the fluorescence intensity continuously increasing from 0-8 h (Fig. 3G). As shown in Fig. 3H, the nanowell loaded with non-CAP/BSA-specific ASC, which lead to no fluorescence (Fig. 3I). Finally, 24 CAP-specific ASCs were retrieved for RmAbs production. The images of single CAP-specific ASCs before picking and after picking are shown in Fig. S3.

**Fig. 3.**
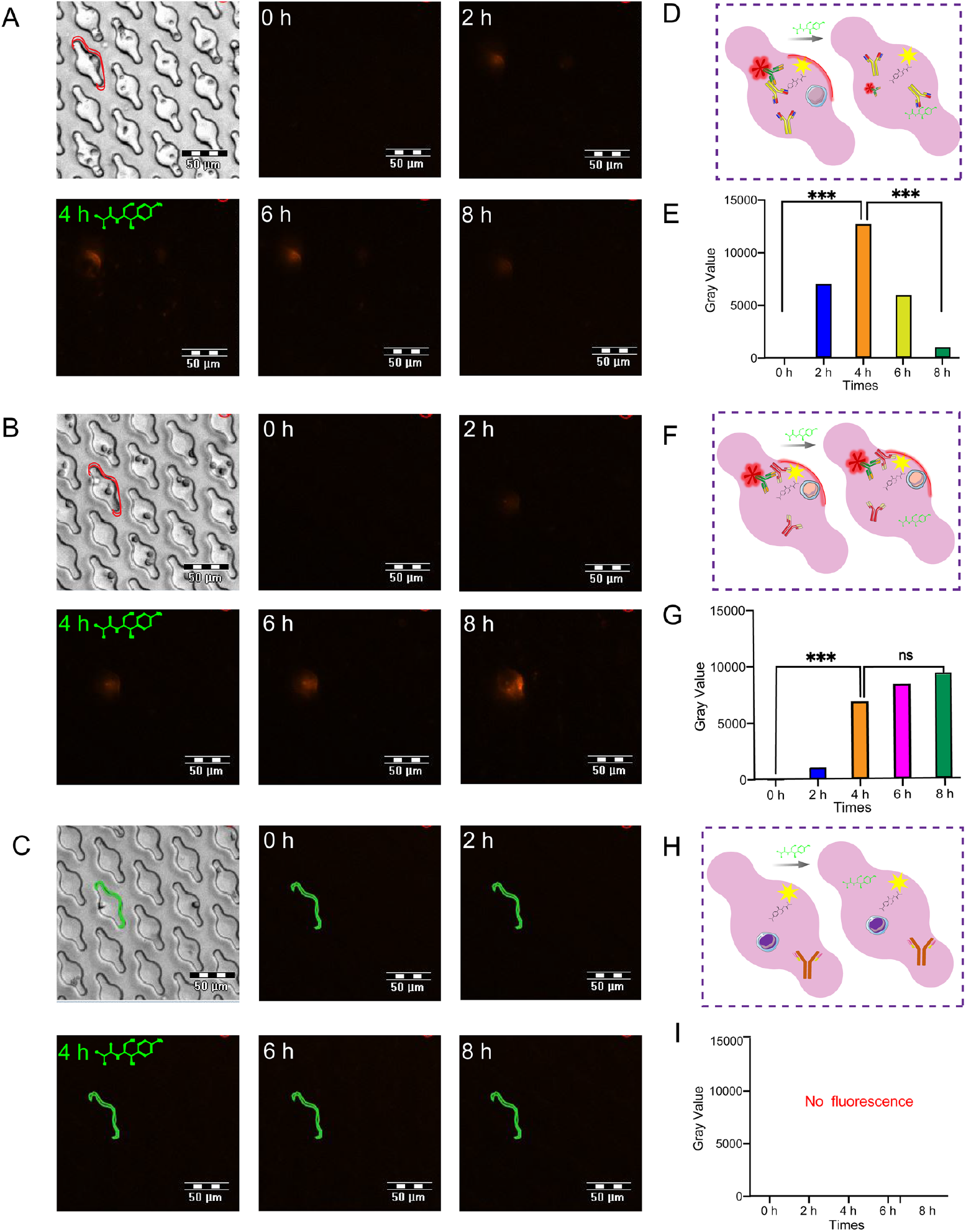
Fluorescence intensity analysis of single ASCs at 0-8 h. (A) Fluorescence intensity of the CAP-specific ASCs at 0-8 h. (B) Fluorescence intensity of the BSA-specific ASCs at 0-8 h. (C) Fuorescence intensity of the non-CAP/BSA-specific ASCs as a control at 0-8 h. (D) Schematic overview of the CSMN for the selection of CAP-specific ASCs. (E) Histogram analysis of fluorescence density of nanowell containing CAP-specific ASC at 0 h-8 h. (F) Schematic overview of the CSMN in the case of nanowell containing BSA-specific ASCs. (G) Histogram analysis of fluorescence density of nanowell containing BSA-specific ASC at 0 h-8 h. (H) Schematic overview of the CSMN in the case of nanowell containing non-CAP/BSA-specific ASCs as a control. (I) The fluorescence density of nanowell containing non-CAP/BSA-specific ASC.

### Characterization of the CAP-specific RmAb

Sixteen pairs of VH and VL were successfully amplified with the forward primer at frame region 1 (FR1) and the reverse primer at the constant region of heavy chain 1 (CH1)/constant region of light chain 1 (CL1) shown in Table S2 and Fig. S4, inserted into expression vectors, and transfected into HEK293T cells for antibody expression. The ELISA and icELISA result of cell supernatant showed that 5 of the 16 yielded RmAbs were positive to CAP-BSA (Fig. S5A) and 2 RmAbs were positive to CAP with the IC_50_ of 4.65 and 0.13 ng mL^-1^, named RmAb2 and RmAb3, respectively (Fig. S5B). The result showed that not all the obtained RmAbs could specifically recognize CAP, which may owe to the production of nonspecific antibodies derived from the accumulation of nonspecific products and PCR errors by extensively amplifying.

Generally, the production of antibodies for diagnosis mainly focuses on the titer, affinity and specificity. However, some antibodies with high affinity and titer in physiological conditions have poor performance in harsh assay environment originally owing to their unstable structure. From a practical viewpoint, the stability of an antibody is more important to develop a robust immunoassay, and an antibody that is able to tolerate physio-chemical factors, such as organic solvents, acid-base and salt, is preferred and highly demanded. Rabbit antibodies possess an additional disulfide bond that is not found in human or mouse antibodies, which theoretically contributes to their higher stability ^32^. Thus, the RmAb3 with higher affinity to CAP, was further purified (Fig. S6) and subsequently assessed for its stability and compared to MmAb for a better application.

As shown in Fig. 4A and 4B, the tolerance of RmAb3 to organic solvents in the icELISA were up to 20% for methanol and 40% for acetonitrile, which were obviously higher than those of MmAb with 10% methanol and only 2.5% acetonitrile (Fig. S7A and S8B). The optimum pH values for RmAb3 and MmAb were comparable, both between 6.5–7.4 (Fig. 4C and S7C). Surprisingly, the RmAb3 showed the best performance in high sodium strength saturated salt solution (Fig. 4D), which was approximately 30-fold higher than that of MmAb with the optimum sodium strength of physiological salt solution (Fig. S7D). Next, the thermal stability and aggregation of the RmAb3 and MmAb were assessed by the Tm and Tagg value, respectively. Fig. 4E shows that the RmAb3 was unfolded at a Tm value of 79.7°C, which was higher than that of MmAb at 74.5°C. Moreover, the Tagg value of the RmAb3 was 81.9°C, and significantly higher than that of MmAb with 74.6°C (Fig. S7E). As shown in Fig. 4F and 4G, the Tm and Tagg values of the RmAb3 were the highest in a saturated salt solution, indicating that the stability of the RmAb3 was highest in the saturated salt solution, whereas the stability of MmAb was the lowest in the saturated salt solution (Fig. S7F and S8G). Wu et al., indicated that the thermal stability related to the binding affinity of antibody^33^. After optimization, the titer and IC_50_ of the RmAb3 were calculated as 24.31 ng mL^-1^ and 0.08 ng mL^-1^, respectively, when the CAP-BSA was used as a coating antigen at the concentration of 37.55 ng mL^-1^ (Fig. 4H), all better than those of the MmAb with 174 ng mL^-1^, 0.28 ng mL^-1^ and 500.0 ng mL^-1^ (Fig. S7H). Furthermore, the stability test proved the RmAb3 was an extremely halophilic protein, which has been rarely reported for antibody. We consider that the outstanding stability and affinity of the RmAb3 in saturated salt may be due to the structural features of the RmAb ^34^. We next analyzed the detailed structure dynamics to explore the halophilic recognition of the RmAb3 and provide insight into the stability of the antibody in harsh environments and useful clues to produce high-quality antibodies for analytical purposes.

**Fig. 4.**
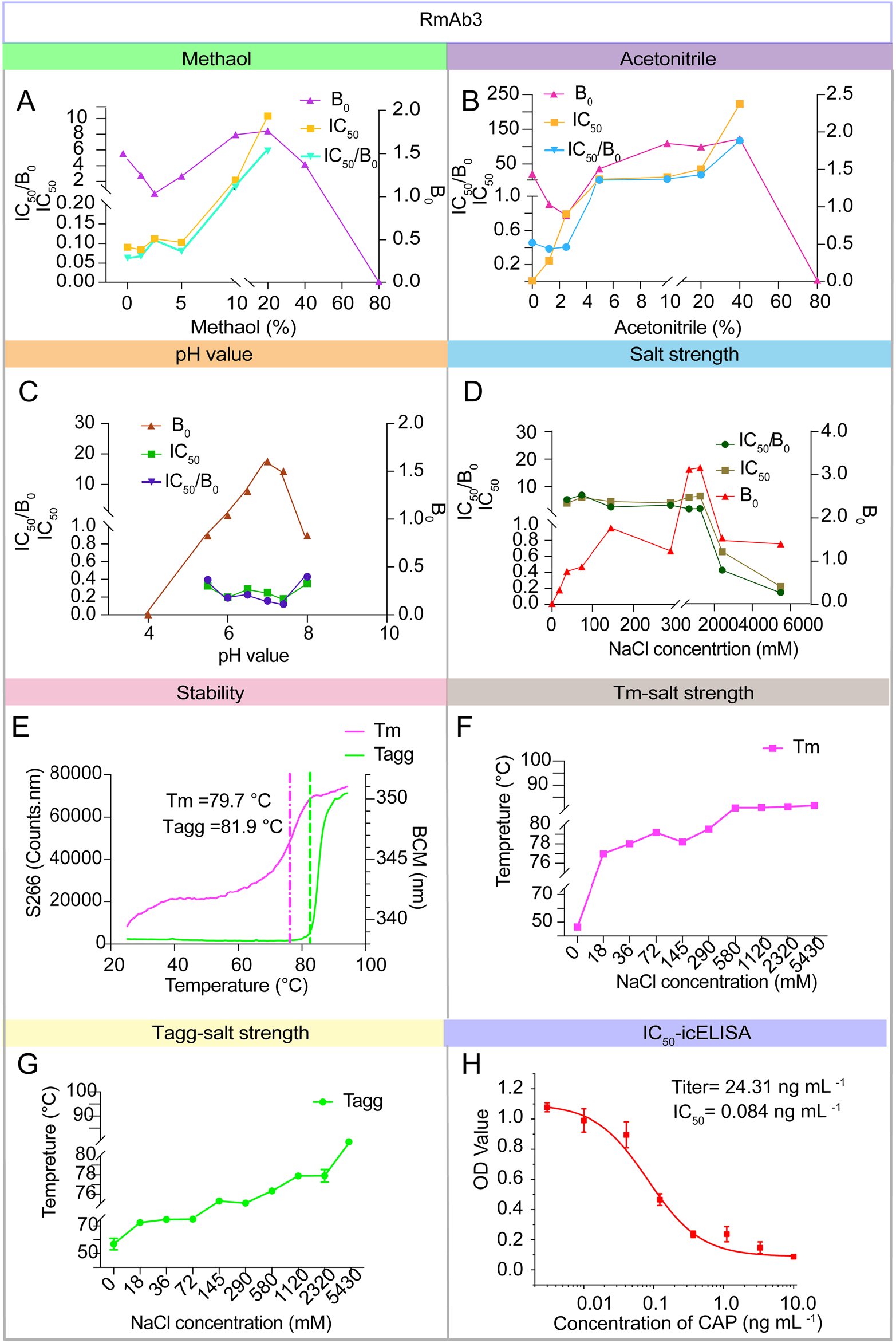
The characteristics of the RmAb3. (A) Methanol tolerance analysis of the RmAb3. (B) Acetonitrile tolerance analysis of the RmAb3. (C) pH tolerance analysis of the RmAb3. (D) Salt strength tolerance analysis of the RmAb3. (E) Thermal stability analysis of the RmAb3. (F) Tm analysis of the RmAb3 at different sodium strengths from 0 M to 5.43 M. (G) Tagg analysis of the RmAb3 at different sodium strengths from 0 M-5.43 M. (H) The affinity analysis of the RmAb3 at the optimum condition in the icELISA.

### Investigation of the halophilic mechanism of the RmAb3

Halophilic proteins are defined as the proteins that can stably work at a certain sodium strength, including slightly halophilic proteins (0.2-0.85 M NaCl), moderately halophilic proteins (0.85-3.4 M NaCl), and extremely halophilic proteins (3.4-5.1 M NaCl) ^35^. For MmAbs, the most suitable working sodium strength is reported to be 0.145 M or below, i.e., non-halophilic proteins ^36, 37^. As shown in the Fig. 4, the RmAb3 was found to be extremely halophilic in a working salt strength of 5.43 M, which has not been reported before for immunoglobulin. Thus, the investigation of how the RmAb3 maintains stable folding and avoids aggregation under high salt conditions is of great significance for its application, and could provide useful information to develop other antibodies with salt tolerance *in vitro*. As the structures of RmAb3 and MmAb show differences including the number of disulfide bonds and the length of the variable region ^34^. Due to their covalent nature, engineered cysteine cross-links have been extensively explored as a means of increasing thermal stability^38^. we hypothesize that the structures of RmAb are critical to the stability of antibody, i.e., extremely halophilic in the study. We studied the molecular mechanisms of the extremely halophilic RmAb3 and the non-halophilic MmAb from the aspects of their structures under control solution, physiological salt solution and saturated salt solution.

We firstly analyzed the detailed the amino acid distribution of the RmAb3-Fv and MmAb-Fv. As shown in Tables S3 and S4, halophilic RmAb3 possessed more negatively charged amino acids of Asp and Glu, and less positively charged amino acids of Lys, Arg and His in the Fv region compared to those of the non-halophilic MmAb. These variations may lead to high density of negative charge on the RmAb3-Fv surface and be positively related to the stability of RmAb in saturated salt solution. We next aligned the VH and VL sequences of the RmAb3 and MmAb. Fig. 5A shows the similarity of the RmAb3 VH and the MmAb VH was 39.52%, and that of RmAb3 VL and MmAb VL was 45.14%, which indicated that the Fv sequences of the RmAb3 and the MmAb were significantly different, especially in the complementarity-determining regions (CDRs). The length of CDRH3 of the MmAb was longer than that of the RmAb3 (17 vs 14 amino acids), while the CDRL3 of the RmAb3 was longer than that of the MmAb (12 vs 9 residues). This observation corresponds to the longer CDRL3 of the rabbit antibody compared to mouse and human antibodies, which is stabilized by the additional disulfide bridges between Cys 81 and Cys 171^34^. The results indicate that the specific amino acid distribution and CDR length of the RmAb3 compared with these of the MmAb would contribute its stability in the harsh environment of saturated salt solution.

**Fig. 5.**
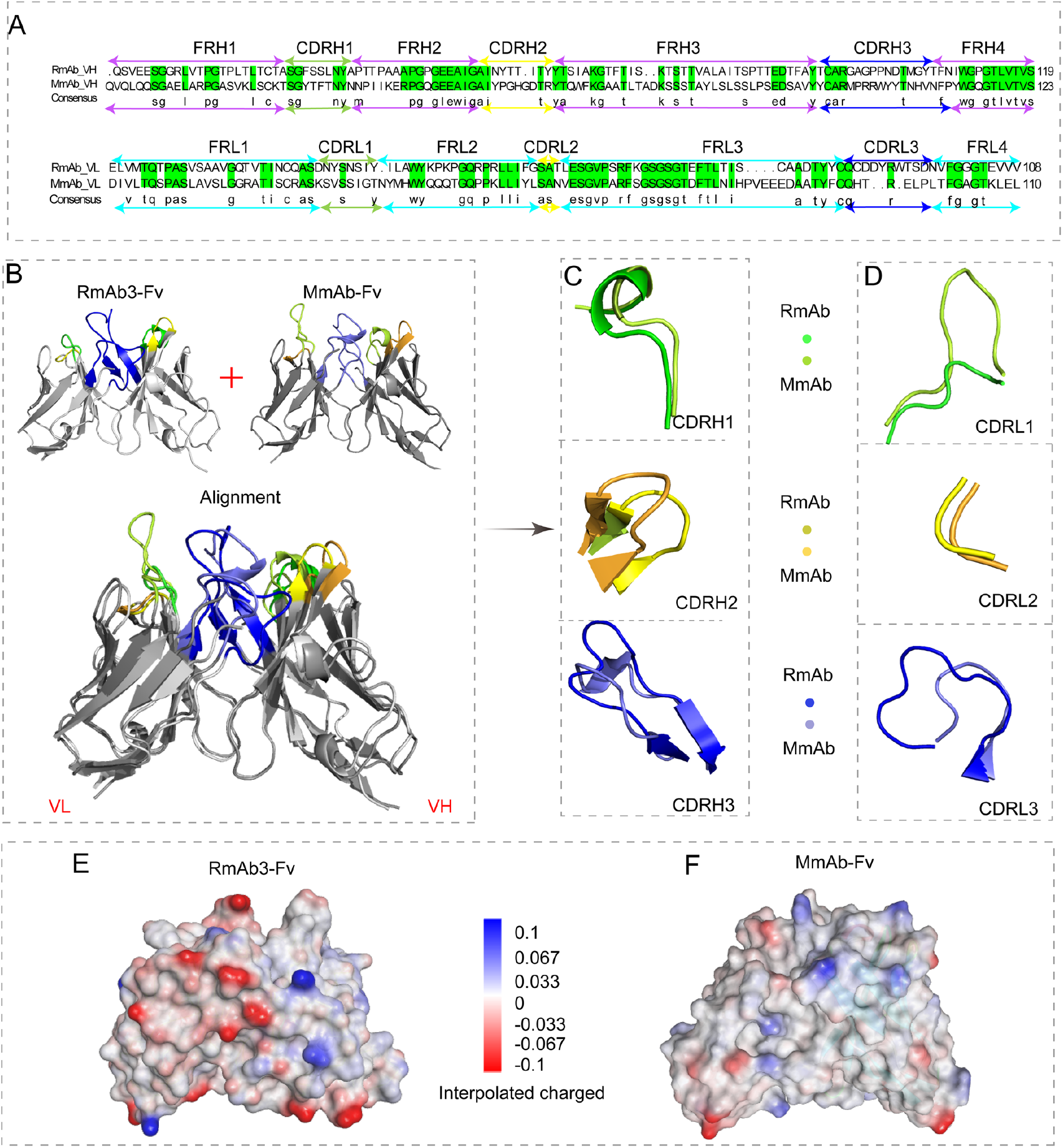
Structural comparison of the RmAb3 and the MmAb. (A) Alignment of the of the RmAb3-Fv and the MmAb-Fv sequences. Amino acids marked by green represent the same amino acid. (B) Alignment of the predicted 3D structures of the RmAb3-Fv and the MmAb-Fv. (C) The detailed alignment of CDRL1, CDRL2 and CDRL3 of the RmAb3-Fv and the MmAb-Fv. (D) The detailed alignment of CDRH1, CDRH2 and CDRH3 of the RmAb3-Fv and the MmAb-Fv. (E) and (F) The charged surface of the RmAb3-Fv and the MmAb-Fv.

To further compare the difference between RmAb and MmAb in the view of two and three dimensions, the structures of the RmAb3-Fv and MmAb-Fv was constructed by the homology modeling. The detailed modeling process of construction and verification were provided in *Supporting Information* (Fig. S8). As shown in Fig. 5B, the structure of RmAb3-Fv was aligned with that of MmAb-Fv and showed an obvious difference, with a root-mean-square deviation (RMSD) of 1.81 Å, especially the loops and helix of the CDRs. The shape of CDRH1, CDRH2 and CDRL2 of the two mAbs was highly similar, whereas that of CDRH3, CDRL1 and CDRL3 was significantly different, as shown in Fig. 5C and 5D. The loop of the hyper-mutated RmAb3 CDRL3 was larger than that of MmAb. The specificity of the RmAb3 CDRs may also contribute to its halophilic characteristics. Then, the surface charges of the RmAb3-Fv and MmAb-Fv were calculated as shown in Fig. 5E and 5F. The surface of RmAb3 was more negatively charged than that of MmAb. It has been reported that Na^+^ interacts with negatively charged side chains on the surface of some proteins, and further stabilizes the structure of the antibody at high salt strength ^39^. In the case of the RmAb3, we speculated that the highly negative surface charge of the RmAb3 renders the antibody more soluble and flexible at high salt strength, contributing to the higher stability. Therefore, the unique amino acid composition especially in the CDR, as well as the secondary structure and high negative surface charge of the RmAb3 all stabilized the antibody structure and then sustained affinity under high salt conditions.

Antibody stability is a prerequisite for binding to antigen under high salt strength assays. The dynamic of antibody-antigen complexes could unveil the allosteric effects in salt solutions ^40^. A total of 100 ns MD simulation was performed to further explore the dynamic of mAbs-CAP complexes, with a time step of 2 fs in control salt solution, physiological salt solution, and saturated salt solution as shown in Fig. S9. The stability and interaction behavior of the RmAb3-CAP and MmAb-CAP complexes were analyzed and compared. Post-simulation analysis, including the average RMSD, solvent accessible surface area (SASA), radius of gyration (Rg), root mean square fluctuation (RMSF), and the number of inter-hydrogen bonds, was used to characterize the stability of mAbs-CAP complexes. The detailed molecular recognition mechanisms of mAbs-CAP in the optimum salt solution were also analyzed to further reveal the halophilic mechanism of the RmAb3.

We observed an increased RMSD value of the RmAb3-CAP complex at 80 s in control salt solution and the MmAb-CAP complex at 90 s in saturated salt solution, indicating that the stability of the RmAb3-CAP complex and MmAb-CAP complex could be influenced by control salt solution and saturated salt solution, respectively. Furthermore, the SASA and Rg of RmAb3-CAP increased in the control salt solution, whereas, those of MmAb-CAP increased in the saturated salt solution. These results of RMSD, SASA, and Rg demonstrated that the stability of mAb-CAP could be significantly damaged in different salt solutions, while the RmAb3-complex was very stable in saturated salt solution, proving to be an extremely halophilic protein (Fig. S10B, S11C, S11B and S12C). Structural flexibility of proteins, as enzymes, has been proven to be essential for their catalytic stability [43, 44]. Therefore, we further characterized the structural flexibility of mAbs-CAP by calculating the RMSF at different sodium strengths. Regarding the RmAb3-CAP complex, the RMSF increased at the positions of 88–98 in the CDRL3 and positions 100–107 in the CDRH3 in the control salt solution. Regarding the MmAb-CAP complex, the RMSF value of positions 97–111 in the CDRH3 increased in the saturated salt solution (Fig. S11D). Additionally, the number of inter-hydrogen bonds between RmAb3 and CAP in saturated salt solution were more than those in the control and physiological salt solutions, as shown in Fig. S10E and S11E. In contrast, the MmAb formed less inter-hydrogen bonds with CAP in saturated salt solutions than in control and physiological salt solutions. These results demonstrate that the RmAb3-CAP was most stable with the highest affinity in a saturated salt solution, further confirming that the RmAb3 is extremely halophilic in a high salt environment.

Finally, the halophilic mechanism of RmAb3 was further verified by analyzing the molecular interaction of mAbs-CAP under different salt strengths. As shown in the top view of the RmAb3-CAP complex in the control salt solution, the nitro of CAP was removed from RmAb3-Fv and transferred to the back of the RmAb3-Fv shown in the Fig. 6A, 6B and 6C (front view), while, the nitro of CAP stably inserts into the binding pocket of the RmAb3 in physiological and saturated salt solutions (Fig. 6D-6I). This indicates that the interactions of RmAb3 and CAP were well stabilized under conditions of high sodium strength, contributing to the high affinity of RmAb to CAP. We also observed that the SASA of the RmAb3 binding pocket was significantly reduced with the increasing sodium strength (Fig. 6J-6L), resulting in increased hydrophobic forces and hydrogen bonds of the RmAb3 and CAP under high salt conditions. In the case of the MmAb, the saturated salt solution led to a conformational change in CAP, which underwent a 180-degree flip, where the nitro of CAP faced the outside of the binding pocket of the MmAb (Fig. 6S-6U). In contrast, the nitro of CAP stably inserted into the binding pocket in control and physiological salt solutions (Fig. 6M-6R), which significantly decreased of the interactions between MmAb and CAP and noticeably impaired their affinity to CAP in saturated salt solution (Fig. 6V-6X).

**Fig. 6.**
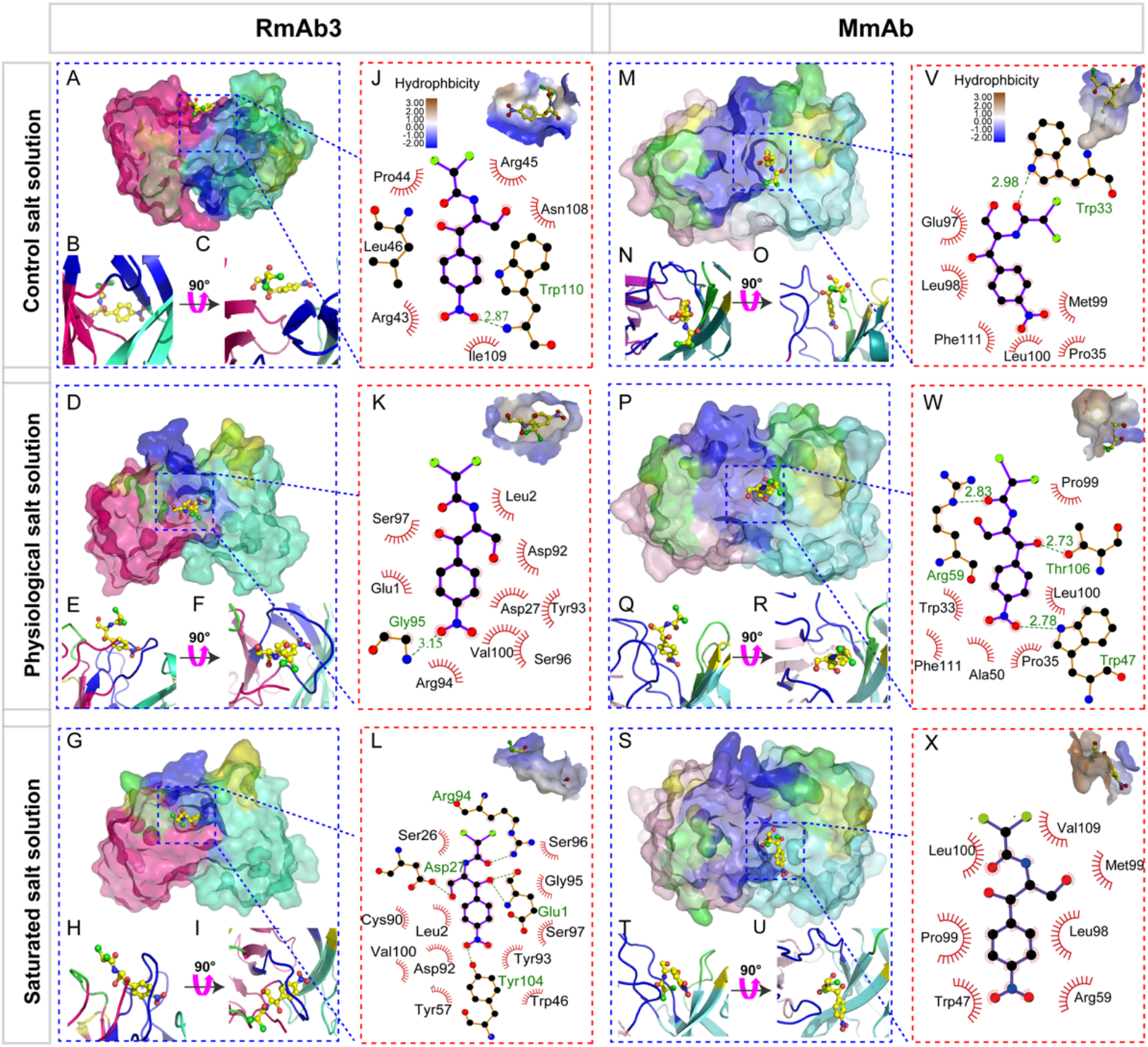
Molecular mechanism analysis of the halophilic RmAb3 and non-halophilic MmAb. (A) and (M) The RmAb3-Fv-CAP complex and MmAb-Fv-CAP complex in control salt solution. (B) and (N) The front view of the RmAb3-Fv-CAP complex and the MmAb-Fv-CAP complex in control salt solution. (N)-(O) The top view of the RmAb3-Fv-CAP complex and MmAb-Fv-CAP complex in control salt solution. (D)-(F) and (P)-(R) The RmAb3-Fv-CAP complex and MmAb-Fv-CAP in physiological salt solution. (G)-(I) and (S)-(U). The RmAb3-Fv-CAP complex and MmAb-Fv-CAP in saturated salt solution. (J) and (V). Hydrophobicity and interaction analysis of the RmAb3-Fv-CAP complex and the MmAb-Fv-CAP in physiological salt solution. The top shows the hydrophobicity analysis of the RmAb3-Fv-CAP complex and MmAb-Fv-CAP binding pocket in control salt solution, the bottom shows the interaction analysis of the RmAb3-Fv-CAP complex and MmAb-Fv-CAP in control salt solution. K and W. Hydrophobicity and interaction analysis of RmAb3-CAP complex and MmAb-CAP in physiological salt solution. (L) and (X). Hydrophobicity and interaction analysis of he RmAb3-Fv-CAP complex and MmAb-Fv-CAP in saturated salt solution. The green dashed line indicates hydrogen bond, and the red arcs indicate hydrophobic interactions.

We next compared the interactions of mAbs-CAP at optimum salt solutions for a better understanding of the RmAb halophilic mechanism, using the RmAb3-CAP in saturated salt solution and the MmAb-CAP in physiological salt solution, respectively. The binding surface area of the RmAb3 is 333.529 Å^2^, which is larger than that of the MmAb (306.716 Å^2^). The large binding pockets of the RmAb3 would relatively provide increased forces and interactions between the antigen and antibody. Furthermore, the orientation and position of CAP in the binding pocket of the RmAb3 and MmAb are also different. Briefly, three hydrogen bonds are formed between Glu1/Asp17/Arg94 and CAP in the binding pocket of RmAb3 (Fig. 6L and 6W), which leads the nitro of CAP towards the “arch” built by the loop of CDRL3 (Fig. 6I), while, another hydrogen bond formed by Tyr104 with CAP drives the CAP into the “arch”. Besides, the hydrophobic force between CAP and the RmAb3 also make a significant contribution to the interactions (Fig. 6L). In the case of the MmAb, three hydrogen bonds are formed between the Trp47/Thr106/Arg59 and CAP, leading the nitro of CAP to vertically insert into the binding pocket (Fig. 6W). The hydrophobic force between the CAP and the MmAb is also important, although it is much weaker than that between the CAP and the RmAb3. Moreover, the interactions between the CAP and the RmAb3 are mainly from the VL, but from the VH in the case of the MmAb-CAP. The observation of the RmAb3 recognition mechanism was corresponding to the previous X-ray crystallography finding that the rabbit light chain and particular CDRL3 was the main contributors to the antibody and antigen interaction^41^. In a word, the binding free energy of RmAb-CAP (−55.84 kcal mol^-1^) in saturated salt solution is obviously lower than that of MmAb-CAP (−41.80 kcal mol^-1^) in physiological salt solution, which contributes to the approximately four-fold higher affinity of the RmAb3 compared to the MmAb.

### Sample evaluation and immunoassay development

Considering the extremely halophilic characteristics, the RmAb3 was used to develop an icELISA for the detection of CAP in biological samples to evaluate the practicality with saturated salt solution as extraction solvent. In the process of sample pretreatment, the samples did not require purification or dilution after extraction, mainly benefiting from the halophilic nature of the RmAb3 and significantly simplifying the assay procedure and reducing the assay time. Whereas the assay based on the MmAb need more labor and time to prepare after extraction. The calculated limits of detection (LODs) of the icELISA based on RmAb3 for CAP were 0.019 µg L^-1^ in milk, 0.02 µg kg^-1^ in pork, and 0.014 µg kg^-1^ in chicken (3 times the standard deviation of the twenty blank samples, 3σ). The accuracy and precision of the developed icELISA were evaluated by recovery and coefficient of variation (CV) in spiked samples. As shown in Table 1, the recoveries of the icELISA with the RmAb3 ranging from 87.9% to 121.2% with intra-assay CVs below 11.7% and inter-assay CVs below 13.7%. To evaluate the reliability of the icELISA, ten practical positive chicken samples confirmed with HPLC-MS/MS were submitted to the developed icELISA, showing a good agreement of two methods (Table S5). Additionally, investigation into the specificity of one immunoassay is also crucial to ensure the accuracy of the subsequent application. As shown in Table S2, the developed icELISA showed ignored cross-reactivities to three analogs of TAP, FF, and FFA below 0.01%. These results indicated that the developed assay based on the RmAb3 with halophilic feature provides high sensitivity, specificity, accuracy and precision with shorter assay time and reduced labor cost, fully meets the requirements of a rapid analytical method for screening purpose.

**Table 1.** Recovery, intra- and inter-assay CV in spiked samples in the icELISA (n=3).

| mAb | Samples | Spiked<br>($\mu\text{g L}^{-1}$ or $\mu\text{g kg}^{-1}$) | Recovery<br>(%) | Intra-assay<br>CV (%) | Inter-assay<br>CV (%) |
| --- | --- | --- | --- | --- | --- |
| RmAb3 | Milk | 0.05 | 96.4 | 7.4 | 9.2 |
|  |  | 0.1 | 87.9 | 9.1 | 10.1 |
|  |  | 0.2 | 107.4 | 10.2 | 13.7 |
|  | Pork | 0.05 | 101.3 | 3.9 | 4.0 |
|  |  | 0.1 | 106.6 | 4.3 | 5.1 |
|  |  | 0.2 | 99.5 | 6.6 | 7.9 |
|  | Chicken | 0.05 | 112.7 | 5.1 | 11.3 |
|  |  | 0.1 | 108.3 | 10.9 | 12.2 |
|  |  | 0.2 | 121.2 | 11.7 | 11.4 |

In this work, a precise RmAb discovery platform named the CSMN, was proposed based on competitive screening single hapten-specific ASC selection in nanowells. CAP is a toxic substance for human health. The CSMN significantly improved the precise of CAP-specific ASC selection and shortened the RmAb preparation time. The titer and affinity of the prepared the RmAb3 is 3-fold higher than that of the best MmAb produced so far. More importantly, the RmAb3 was found to be extremely halophilic, which has not been reported previously. We found that the unique structure, negatively charged surface, and reduced SASA of the binding pocket of the RmAb3 all contributed the extremely halophile. This novel CSMN for RmAb discovery and findings of antibody halophilic mechanism in the study provide a general and efficient tool for the development of new generation of immunoassay for small molecule in various fields.

## Materials and methods

### Materials and apparatus

The chloramphenicol (CAP), hemocyanin-keyhole limpet (KLH), bovine serum albumin (BSA), red blood cell lysis buffer, FRMI 1640 medium, fetal bovine serum (FBS), and SuperScript™ III CellsDirect™ cDNA synthesis kit were supplied by Thermo Fisher Scientific, Inc. (Waltham, MA, USA). The rabbit spleen lymphocyte separation kit and carbonate buffer solution (CBS) were obtained from Solarbio Life Sciences, Inc. (Beijing, China). The horseradish peroxidase (HRP)-labeled goat anti-rabbit IgG and phycoerythrin (PE)-anti-rabbit F(ab’)_2_ antibody was obtained from Jackson ImmunoResearch Laboratories, Inc. (West Grove, PA, USA).

White polystyrene micro-titer plates and syringe filters (0.45 µm) were obtained from Costar, lnc. (Milpitas, CA, USA). H100 hexagonal nanowell (size of 100 µm, depth of 100 µm, 60,000, 0.9 nL/nanowell), U40 UFO nanowell (size of 40 µm, depth of 40 µm, 50,000 wells, 50 pL/nanowell), U25 UFO nanowell (size of 25 µm, depth of 25 µm, 20,000 wells, 12 pL/nanowell) and 370K nanowell (size of 20 µm, depth of 25 µm, 370,000 wells, 8.6 pL/nanowell) were obtained from Automated-Lab-solutions GmbH. Inc (Jena, Germany). Specific pathogen-free, female New Zealand white rabbits were used in all experiments and were purchased from Beijing Vital River Laboratory Animal Technology Company (Beijing, China).

The optical density (OD) value was measured via the PerkinElmer Envision plate reader (Waltham, MA, USA). Count star was used to count the number of splenocytes (Shanghai, China). single ASCs were selected by the CellCelector™ (Jena, Germany). Melting temperature (Tm) and aggregation temperature (Tagg) of the mAbs were measured by the Unit High-throughput Protein Stability Analyzer (Pleasanton, CA, USA).

### Rabbit immunization and cell preparation

All animal procedures of six 2-month-old female New Zealand white rabbits were approved by the Animal Ethics Committee of China Agricultural University and strictly conducted in accordance with Chinese laws and guidelines. The immunogen (CAP-KLH), which had been prepared in our previous work ^42^, was dissolved in 0.6 mL of PBS emulsified with Freund’s adjuvant for the first immunization, and Freund’s incomplete adjuvant for further boosting immunization. Then, the water-in-oil mixture was subcutaneously injected into rabbits at multiple sites. Each was injected six times at 4-week intervals. Antisera was collected from the ear veins after the injection and assayed by icELISA with the coating of CAP-BSA, the icELISA protocol is shown in the *Supporting Information*.

The spleen of the rabbit with the highest antisera affinity was taken under sterile conditions. The cells were blown out of the spleen with 10 mL of the FRMI 1640 with 5% FBS, cell suspension was filtered through a 45 µm cell strainer, centrifuged for 10 min at 1000 rpm, resuspended in RPMI 1640 and transferred to a 15 mL of conical tube. Then, 1mL of the red blood cell lysis buffer was added to the tube, mixed gently, and incubated at room temperature for 15 min, Finally, the cell suspension was filtered through a 45-µm cell strainer and adjusted to a concentration of 1 × 10^9^ cells mL^-1^.

### Nanowell optimization and surface modification

To test the efficiency of the nanowell for single-cell occupancy, we have optimized a range of nanowells, including a hexagonal nanowell (H100), UFO nanowell (U40), UFO nanowell (U25) and SIEVEWELL nanowell (370K), with nanowell sizes ranging from 100 µm to 20 µm and nanowell volumes ranging from 0.9 nL to 8.6 pL. The single-cell occupancy was calculated according to Eq. 1, with the number of plated rabbit-splenocytes half of the nanowell numbers in the chip.

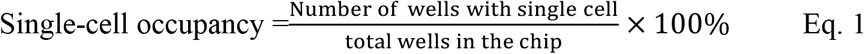

The optimal nanowell with the highest single-cell occupancy was pretreated with 1 mL of anhydrous ethanol and centrifuged at 1000 rpm for 10 min to drain the bubbles. Next, 0.5 mL of anhydrous ethanol was removed, and the chip was washed five times with CBS (0.05 M, pH = 9). The nanowell was coated with 0.5 mL of CAP-BSA (1 µg mL^-1^) at 4°C for 16 h and blocked with 0.5 mL of 2% BSA at 37 °C for 1 h.

### CSMN procedure development

The number of the rabbit splenocytes loaded in the optimal nanowell was further optimized. The splenocytes were incubated with the PE-anti-rabbit F(ab’)_2_ antibody and evenly plated into the nanowells by a pipettor in a “Z”-shape. The nanowell was centrifuged at 1500 rpm for 3 min with slow deacceleration, and primed with RPMI 1640 medium. The splenocytes in the nanowells were scanned and imaged under a microscope with autofocus capability at both brightfield and fluorescence field in the CellCelector™ platform. The nanowell was incubated at 37°C with 5% CO_2_ to enable antibody secrete from the loaded ASCs, bind to the CAP-BSA coated in the nanowell and aggregate PE-anti-rabbit F(ab’)_2_ antibody. Then the fluorescence intensity of nanowell was scanned every 2 h to monitor the antibody secretion from the loaded ACS for 8 h. At h 4, the CAP (1 µg mL^-1^) was added in the nanowell, as a competitor of the coated CAP-BSA for the secreted CAP-specific antibody from the single ASC. The ASCs with the fluorescence intensity reaching the peak at h 4 and decreasing with the addition of CAP at 4-8 h, were selected as the positive CAP-specific ASCs; these ASCs were automatedly picked by a 20-µm diameter glass capillary in a robotic arm in the CellCelector™ platform. Then, the CAP-specific ASC was transferred into a 0.2-mL tube containing 10 µL of lysis buffer and 1 µL of RNaseOUT^™^ (40 U µL^-1^) for subsequent antibody gene production (*Supporting Information*). All data were analyzed using statistical software (SPSS 21.0). One-way analysis of variance was used for comparisons among multiple groups. Tukey’s post hoc test was used for pairwise comparisons of the mean values between multiple groups.

### CAP-specific RmAb production and characterization

The variable region of the heavy chain (VH) and variable region of the light chain (VL) of the RmAb were amplified by polymerase chain reaction (PCR) with the specific primers. The sequences of the paired VH and VL of single ASCs were analyzed by the TA cloning and Sanger sequencing; the detailed process is shown in *Supporting Information*. According to the preference of the human cell expression system, the codon of the amplified VH and VL genes were optimized, and then the whole gene of the variable region was synthesized and cloned directly into the mammalian cell expression vector pFUSE-rabbit Fc, which contained the Fc constant region of the rabbit antibody. The plasmid was transfected into the HEK293 cells (*Supporting Information*), and the supernatant was collected and centrifuged for 5 min to remove the cells. The RmAb was purified by Protein A affinity column.

### CAP-specific RmAb3 characterization

The CAP-specific RmAb3 from the ASCs were characterized and compared to the CAP-specific MmAbs by antibody titer, affinity, specificity, stability and tolerance of sodium strength, methanol, acetonitrile, and pH. All measurements were conducted for three replicates.

The antibody titer is represented by antibody dilution, and the antibody affinity is represented by the half maximal inhibitory concentration (IC_50_) values from the standard curves of the icELISA for CAP based on mAbs. The standard curves of the icELISA were constructed by OriginPro 8.0 (OriginLab Corp., Northampton, MA) and data were fitted to the following four-parameter logistic equation according to the Eq. 2.

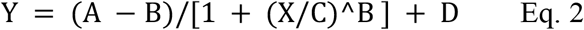

Where A represents the responses at high asymptotes of the curve, B acts as the slope factor, C is the IC_50_ of the curve, D is the responses at low asymptotes of the curve, and X is the calibration concentration.

The specificity of RmAb3 was evaluated by using thiamphenicol (TAP), florfenicol (FF) and florfenicolamide (FFA). The cross reactivity (CR) was calculated according to the Eq. 3:

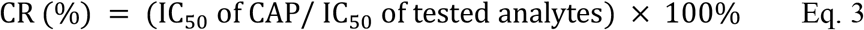

The stability of the mAbs was assessed by Tm and Tagg. To determine the onset of aggregation, a thermal ramp between 25°C and 100°C was used with a heating rate of 1°C per minute. Tm values were calculated from the fluorescence data in terms of the barycentric mean (BCM), while Tagg values were calculated based on the 266 nm static light scattering (SLS).

The tolerances of sodium strength, methanol, acetonitrile, and pH were assessed by the IC_50_/B_0_ in the icELISA. The B_0_ is the OD value of the icELISA in the absence of CAP.

### RmAb3 halophilic mechanism investigation

The three-dimensional structures of the RmAb3 and MmAb fragment variable region (Fv) prepared in our laboratory were constructed by homology modeling in the Discovery Studio 2019 software, and docking analysis was performed based on a grid-based semi-flexible molecular docking software of CDOCKER. The detailed processes of the homology modeling and docking are shown in *Supporting Information*. The best docking model with the lowest CDOCKER energy and higher fitness value was refined with a 100 ns molecular dynamic (MD) simulation by GROMACS5.0 program software ^43^ (*Supporting Information*). To mimic the conditions of different concentrations of NaCl, three simulation systems were prepared for RmAb3-Fv-CAP and MmAb-Fv-CAP, respectively, containing 0 M NaCl (control salt solution), 0.145 M NaCl (physiological salt solution), and 5.43 M NaCl (saturated salt solution) by GROMASC5.0 in the KBFF force field ^44^ with the SPC/E solvent model ^45^. The solvated structures were minimized by the steepest descent method for 15,000 steps at 310 K temperature and constant pressure. The LINCS algorithm was used to constrain the bond length ^46^, the electrostatics interactions were calculated using a PME algorithm^47^, and the binding free energy of mAbs and CAP was calculated using the MM/PBSA method after the MD process^48^.

### Sample evaluation and immunoassay development

The practicability of the developed icELISA based on the RmAb3 with high affinity and stability was evaluated by spiked biological samples and practical CAP-positive chickens that were already confirmed by high performance liquid chromatography-tandem mass spectrometry (HPLC-MS/MS) as in detailed described in the *Supporting Information*. The CAP-negative samples of milk, pork, chicken, and practical CAP-positive chickens were acquired from the Beijing Key Laboratory of Diagnostic and Traceability Technologies for Food poisoning (Beijing, China).

## Notes

The authors declare no competing financial interests.

## Acknowledgments

This work was supported by the National Science Foundation of China (Grant No. 32172905, 32172910), the National Key R&D Program of China (2018YFC1602900) and the Sanming Project of Medicine in Shenzhen (SZSM201611068).

