## Supporting method, table and figure for "Rare Monoclonal Antibody Discovery Based on Indirect Competitive Screening of Single Hapten-specific Rabbit Antibody Secreting Cell"

### Contents

1. Methods

1.1 Regents and apparatus

1.2 Characterization of rabbit antisera

1.3 Optimization of nanowell

1.4 Fluorescence intensity of the U25

1.5 Preparation of single ACS antibody gene

1.6 Production of single ACS antibody

1.7 Characterization of the antibody from the isolated ACSs

1.8 Characterization of the CAP-specific MmAb

1.9 Homology modeling and molecular docking of the RmAb3-Fv-CAP

1.10 MD of the RmAb3-Fv-CAP and the MmAb-Fv-CAP

1.11 Sample evaluation and immunoassay development

2 Results

2.1 Characterization of rabbit antisera

2.2 Optimization of nanowell

2.3 Fluorescence intensity of the U25

2.4 Preparation of single ACS antibody gene

2.5 Production of single ACS antibody

2.6 Characterization of the antibody from the isolated ACSs

2.7 Characterization of the CAP-specific MmAb

2.8 Homology modeling and molecular docking of the RmAb3-Fv-CAP

2.9 MD of the RmAb3-Fv-CAP and the MmAb-Fv-CAP

2.10 Sample evaluation and immunoassay development

#### 1. Methods

##### 1.1 Reagents and apparatus

SuperScript™ III CellsDirect™ cDNA synthesis kit, Opti-MEM and Lipofectamine 2000 were purchased from Thermo Fisher Scientific, Inc. (Waltham, MA, USA). Gel recovery kit and Taq PCR Master Mix were obtained from Takara Biomedical Technology Co., Ltd, Inc. (Kyoto, Japan). 3,3',5,5'-Tetramethylbenzidine (TMB) substrate were purchased from J&K Chemical company (Shanghai, China). Luria-Bertani (LB) medium was purchased from Solarbio life science (Beijing, China). Chloramphenicol (CAP), keyhole limpet hemocyanin (KLH) and bovine serum albumin (BSA) were supplied by Sigma-Aldrich (St. Louis, MO, USA). The rabbit spleen lymphocyte separation kit and carbonate buffer solution (CBS) were obtained from Solarbio Life Sciences, Inc. (Beijing, China). The horseradish peroxidase (HRP)-labeled goat anti-rabbit IgG and phycoerythrin (PE)-anti-rabbit F(ab')$_2$ antibody were obtained from Jackson ImmunoResearch Laboratories, Inc. (West Grove, PA, USA). White polystyrene micro-titer plates and syringe filters (0.45 μm) were obtained from Costar, lnc. (Milpitas, CA, USA).

The optical density (OD) value was measured via the PerkinElmer Envision plate reader (Waltham, MA, USA). Polymerase chain reaction (PCR) was performed in the Applied Biosystems PCR Thermal Cycler (Waltham, MA, USA). Fluorescence intensity was analyzed by the software in the CellCelector™ platform (Jena, Germany).

The following buffers were used in the enzyme-linked immunosorbent assay (ELISA) and indirect competitive enzyme-linked immunosorbent assay (icELISA) procedure: coating buffer, 0.05 M carbonate buffer, pH 9.6; blocking buffer: sodium phosphate-buffered saline (PBS) with 0.5% casein, containing 1% BSA and 0.1% Proclin-300, pH 7.4; PBS buffer (0.01 M, pH 7.4); washing buffer (0.01 M PBS, 0.05% Tween 20, pH 7.0); stop solution (2 M $H_2SO_4$).

1.2 Characteristic of rabbit antisera and preparation of splenocytes

Antisera was collected from the ear veins of immunized rabbit after the sixth injection and assayed by icELISA. The icELISA protocol were carried out as follows: coating antigens (100 μL well$^{-1}$) diluted with carbonate buffer were added to a microplate, and the microplate was then placed in an incubator at 37°C for 2 h. After washing the microplate three times using washing buffer, the microplate was blocked using blocking buffer at 37°C for 1 h. Then the blocking buffer was discarded. 50 μL of PBS and 50 μL of diluted antisera was sequentially added to the microplate (ELISA); or 50 μL of standard solution (diluted CAP or other analytes) and 50 μL of diluted antisera was sequentially added to the microplate (icELISA). After incubation at 37°C for 30 min, the microplate was washed three times, followed by the addition of HRP-labeled goat anti-rabbit IgG secondary antibody (1:5000, 100 μLwell$^{-1}$). Next, the microplate was washed four times, and then TMB substrate (100 μL well$^{-1}$) was added to the microplate, followed by incubation for 15 min at 37°C. Then, stop solution (50 μL well$^{-1}$) was added to the microplate. Finally, the OD values were measured at 450 nm.

The cells were blown out of the spleen with 10 mL of the FRMI 1640 with 5%

FBS, cell suspension was filtered through a 45 μm cell strainer, centrifuged for 10 min at 1000 rpm, resuspended in RPMI 1640 and transferred to a 15 mL of conical tube. Then, 1mL of the red blood cell lysis buffer was added to the tube, mixed gently, and incubated at room temperature for 15 min, Finally, the cell suspension was filtered through a 45-μm cell strainer and adjusted to a concentration of $1 \times 10^9$ cells $mL^{-1}$.

1.3 Optimization of nanowell

Four nanowells of H100, U40, U25 and 370K, and were chosen to detect the optimum chip for splenocytes seeding and selection. To test the single-cell occupancy of the nanowell, the number of loaded cells was adjusted to the half of the well numbers i.e., $3 \times 10^4$ cells $mL^{-1}$ for 100-μm-diameter H100, $2.5 \times 10^4$ cells $mL^{-1}$ for 40-μm-diameter U40, $1 \times 10^4$ cells $mL^{-1}$ for 25-μm-diameter U25, and $1.5 \times 10^5$ cells $mL^{-1}$ for 20-μm-diameter 370K.

1.4 Fluorescence intensity of U25

The fluorescence intensity was analyzed based on the automated inverted fluorescence microscope with a high-speed scanning stage in the CellCelector™ platform. The gray values of five regions represented the U25 were analyzed by the software in the CellCelector™ platform.

1.5 Preparation of single ACS antibody gene

After single B cell isolation, the PCR tube containing the single CAP-specific ACS was thawed at 70°C for 20 min. Then 5 μL of DNase I (1 U $\mu L^{-1}$) and 2 μL of 10 × DNase I Buffer was added into the tube to degrade the DNA. 1 μL of EDTA (25 mM) was added to the tube and incubated at 70°C for 15 min to inhibit the activity of DNase I. Then the PT-PCR of the single CAP-specific ACS RNA was performed with 2 μL of Oligo(dT)$_{20}$ (50 mM) and 1 μL of 10 mM dNTP Mix I, incubated at 70°C for 5 min, placed on ice for 2 min and added 6 μL of 5 × RT Buffer, 1 μL of RNaseOUT™ (40 U $\mu L^{-1}$), 1 μL of SuperScript™ III RT (200 U $\mu L^{-1}$) and 1 μL of 0.1 M DTT, the tube was transferred into a thermal cycler preheated to 50°C and incubated for 50 minutes, and the reaction was inactivated at 85°C for 5 min. 1 μL of RNase H (2 U $\mu L^{-1}$) was added to each tube and incubated at 37°C for 20 min. Finally, the DNA of the single ACSs were obtained and the variable region genes of the antibody were amplificated by PCR with the specific primers.

The components of the PCR reaction system to amplify variable region of heavy chain (VH) and variable region of light chain (VL) genes were as follows: 1 μL of single ASC cDNA, 10 μL of PCR SuperMix, forward primer and sense primer (0.5 μM). The PCR was run using the following program: 5 min at 94°C; 25 cycles of 30s at 94°C, 45s at 55°C, 45s at 72°C; and 7 min at 72°C. The DNA products are separated using agarose gel electrophoresis. Briefly, 5 μL of PCR product is mixed with 1 μL of DNA loading buffer and loaded into 1% (w/v) agarose gel, the gel was imaged by using UV light assisted visualization of bands at about 400 bp. The target bands were cut and recovered. Then the product was purified by the PCR product purification kit and linked to the pMD18 T vector overnight at 16°C. The conjugates were transferred into competent cells DH5α and cultured in antibiotic-free LB medium at 37°C for 1 h. The conjugates were coated on ampicillin-containing LB solid medium overnight. The next day, the monoclonal colony in good condition was selected for shaking culture for 3 to 5 h, and then the liquid was used for Sanger sequencing.

1.6 Production of single ACS antibody

According to the preference of human cell expression system, the codon of the amplified heavy and light chain genes was optimized, and then the whole gene of variable region was synthesized and cloned directly into the commercial mammalian cell expression vector pFUSE-rabbit Fc, which contained the Fc constant region of rabbit antibody. HEK293 cells were counted, paved and cultured overnight at 37°C and 5% $CO_2$. The cells with the confluence of 70%–80% was ready for transfection, then the plasmid was diluted with a certain volume of Opti-MEM, carefully mixed, and labeled as liquid A. The transfection reagent Lipofectamine 2000 was diluted with the same volume of Opti-MEM, named liquid B. The mass ratio of plasmid to PEI was 1: 2 and kept at room temperature for 15 min. Liquid B was slowly added to liquid A and mixed. It was kept at room temperature for 20 min and slowly added to the cell culture medium. The supernatant was collected and centrifuged for 5 min to remove the cells after 48 h of transfection. The antibody was purified by Protein A affinity column.

1.7 Characterization of the antibody from the isolated ACSs

The affinity of the 16 RmAbs from the CAP-specific ACSs were evaluated by the ELISA and icELISA. Non-denaturing gel electrophoresis and denaturing gel electrophoresis were performed to evaluate the the RmAb3.

The antibody titer is represented by antibody dilution, and the antibody affinity is represented by the half maximal inhibitory concentration ($IC_{50}$) values from the standard curves of the icELISA for CAP based on mAbs. The standard curves of the icELISA were constructed by OriginPro 8.0 (OriginLab Corp., Northampton, MA) and data were fitted to the following four-parameter logistic equation according to the Eq. S1.

$$Y = (A - B)/[1 + (X/C)^B] + D \qquad \text{Eq. S1}$$

Where A represents the responses at high asymptotes of the curve, B acts as the slope factor, C is the $IC_{50}$ of the curve, D is the responses at low asymptotes of the curve, and X is the calibration concentration.

The specificity of RmAb3 was evaluated by using thiamphenicol (TAP), florfenicol (FF) and florfenicolamide (FFA). The cross reactivity (CR) was calculated according to the Eq. S2:

$$CR\,(\%) = (IC_{50} \text{ of CAP}/\ IC_{50} \text{ of tested analytes}) \times 100\% \qquad \text{Eq. S2}$$

The stability of the mAbs was assessed by Tm and Tagg. To determine the onset of aggregation, a thermal ramp between 25°C and 100°C was used with a heating rate of 1°C per minute. Tm values were calculated from the fluorescence data in terms of the barycentric mean (BCM), while Tagg values were calculated based on the 266 nm static light scattering (SLS).

The tolerances of sodium strength, methanol, acetonitrile, and pH were assessed by the $IC_{50}/B_0$ in the icELISA. The $B_0$ is the OD value of the icELISA in the absence of CAP.

1.8 Characterization of the CAP-specific MmAb

The affinity of the CAP-specific MmAb was evaluated by the ELISA and icELISA. The stability of CAP-specific MmAb was evaluated by Tm and Tagg. Optimization of the sodium strength of the icELISA based on the CAP-specific MmAb was performed to characterize the sodium tolerance of MmAb. Besides the stability of CAP-specific MmAb at different concentration of sodium was evaluated by Tm and Tagg.

1.9 Homology modeling and molecular docking of the RmAb3-Fv-CAP

The three-dimensional (3D) structures of the fragment variable region (Fv) of the RmAb3-Fv was constructed by Discovery Studio 2019 (DS2019) software (Dassault Syatèmes BIOVIA, San Diego, CA). Template structures of VH and VL were first identified by BLAST search in the PDB database. Five antibody crystal structures were selected as the templates. The structure of the Fv was then constructed by superimposing these templates to determine the relative spatial orientation of the heavy and light chains. The highest quality model (with the lowest probability density function energy and highest discrete optimized protein energy) was selected to optimize the complementarity-determining regions (CDRs) by aligning the published crystal structures through the IMGT/V-QUEST database (http://www.imgt.org).

Ramachandran plots and Profile-3D analysis were applied to evaluate the resultant homology model.

Docking analysis was performed to investigate the specific binding mechanism of the RmAb3-Fv-CAP by CDOCKER, a grid-based semiflexible molecular docking method in the DS2019 program. For the Fv, a single docking pose in the Complementarity-determining region (CDR) region was found based on the grid spacing and minimum site size of the antibody binding pocket. The pre-optimized small molecular model of CAP was then docked into the cavity of the binding pocket formed by the CDR regions.

1.10 MD of the RmAb3-Fv-CAP and the MmAb-Fv-CAP

The Molecular dynamics simulation (MD) of the RmAb3-CAP complex and MmAb-CAP complex was performed in three concentration of NaCl to demonstrate the halophilic mechanism of the the RmAb3 and non-halophilic mechanism of MmAb by GROMASC5.0 in the KBFF force field (Weerasinghe et al., 2003) with the SPC/E solvent model (Jorgensen et al., 1983). The solvated structures were minimized by the steepest descent method for 15,000 steps at 310 K temperature and constant pressure. The LINCS algorithm was used to constrain the bond length (Hess et al., 1997), the electrostatics interactions were calculated using a PME algorithm (Darden et al., 1993),

1.11 Sample evaluation and immunoassay development

Under the optimum conditions, the sensitivity and specificity of icELISA were explored using $IC_{50}$ and limits of detection (LOD). The LOD was determined based on 20 blank samples of milk, pork and chicken and was calculated by the average value plus three times the standard deviation. The recovery test was used to evaluate the accuracy of the icELISA. Briefly, blank samples were spiked with CAP at three different concentrations, and the spiked samples were then submitted to the icELISA for recovery analysis after pretreatment. The detailed preparation of the samples is presented as follows: milk is without preparation; 3.0 g of pork was homogenized and mixed with 6 mL of ethyl acetate and hexane in a 50 mL tube. After vertexing for 10 min, the mixture was centrifuged at 5000 rpm for 10 min, 4 mL of supernatant was separated and dried by nitrogen at 60°C. The residue was dissolved in 3mL of PBS buffer and could be used for analysis; 3.0 g of chicken was homogenized and mixed with 6 mL of ethyl acetate and hexane in a 50 mL tube. After vertexing for 10 min, the mixture was centrifuged at 5000 rpm for 10 min, 4 mL of supernatant was separated and dried by nitrogen at 60°C. The residue was dissolved in 3 mL of PBS buffer, and after filtering through a filter membrane (0.22 μm), the sample solution could be used for analysis. The CAP-negative samples of milk, pork, chicken, and practical CAP-positive chickens were acquired from the Beijing Key Laboratory of Diagnostic and Traceability Technologies for Food poisoning (Beijing, China).

The high-performance liquid chromatography-tandem mass spectrometry (HPLC-MS/MS) method for the CAP detection in 12 positive chicken was as follows: A Varian 1200 L triple-quadrupole tandem mass spectrometer (Palo Alto, CA) coupled with a ProStar 410 autosampler and two ProStar 210 pumps and a 1200 L triple-quadrupole mass spectrometer were used with an ESI source. The Varian MS workstation version 6.7 software was used for data acquisition and processing. Chromatographic separation was performed on a Zorbax Column Eclipse XDB C8 (4.6 mm × 150 mm i.d., 3 μm) (Milford, MA). The mobile phase consisted of (A) acetonitrile 80% (v/v) and (B) double distilled water 20% (v/v) containing 0.1% formic acid. The mobile phase, previously degassed with high-purity helium, was pumped at a flow rate of 0.4 mL/min, and the injection volume was 10 μL. ESI was operated in the positive and negative ion mode. The electrospray capillary potential was set to 65 V, the needle at 5850 V, and the shield at 750 V. Nitrogen at 48 mTorr and 375°C was used as a drying gas for solvent evaporation. APCI was operated in the positive mode. The capillary potential was set to 65 V, the APCI torch at 450°C, and the shield at 750 V. Nitrogen at 48 mTorr was set at 400°C. Full-scan spectra were obtained in the ranges of 250-800 amu for CAP detector at 1450 V. For both ESI and APCI the atmospheric pressure ionization (API) housing was kept at 50°C. Parent compounds were subjected to collision-induced dissociation using argon at 3.80 mTorr in the multiple reaction monitoring (MRM) positive and negative mode. The scan time was 1 s, and the detector multiplier voltage was set to 1450 V, with an isolation width of m/z 1.2 for quadrupole 1 and m/z 2.0 for quadrupole 3.

#### 2 Results

##### 2.1 Characterization of rabbit antisera

The $IC_{50}$ represents the antisera affinity and is shown in Table S1.

Table S1. Characterization of rabbit antisera.

| Antisera[a] | CAP-BSA[b] |
| --- | --- |
| | $IC_{50}$[c] |
| rabbit # 1 | 0.31 ng $mL^{-1}$ |
| rabbit # 2 | 1.20 ng $mL^{-1}$ |
| rabbit # 3 | 0.88 ng $mL^{-1}$ |
| rabbit # 4 | 1.05 ng $mL^{-1}$ |
| rabbit # 5 | 3.22 ng $mL^{-1}$ |
| rabbit # 6 | 0.27 ng $mL^{-1}$ |

a Antisera were diluted 20,000-fold with the coating of CAP-BSA. b The concentrations of CAP-BSA was 10 ng $mL^{-1}$. c The $IC_{50}$ analysis of six rabbit according to Eq. 2.

#### 2.2 Optimization of nanowell

Optimization of the four nanowells of H100, U40, U25 and 370K, and is aimed to find the optimum nanowell for splenocytes with the highest single-cell occupancy. As shown in Figure S1, the U25 was chosen as the optimum chip for splenocytes. Then the splenocyte number plated in the U25 was further optimized to presume higher single-cell occupancy.

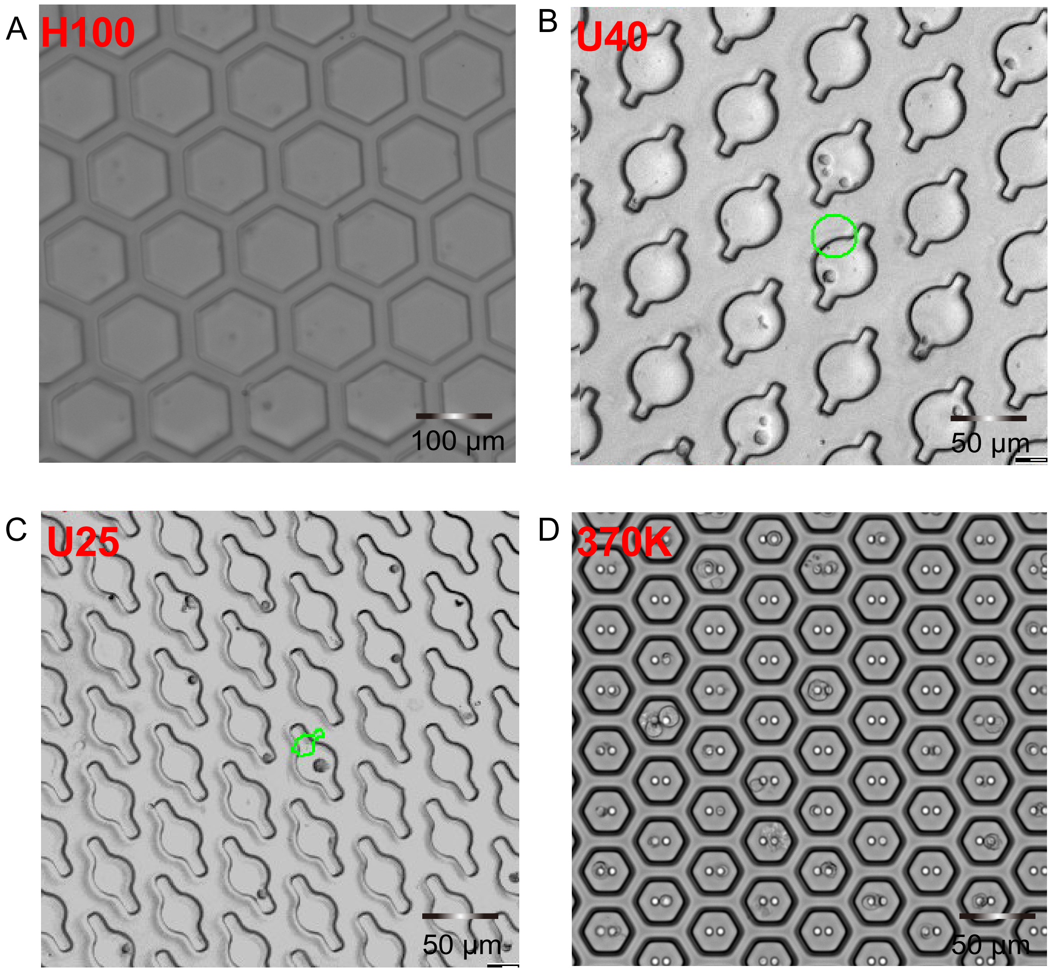


Alt Text: Four images of nanowells with different diameter size.

Figure S1. The optimization of the nanowell diameter size. A: H100 with 100-μm-diameter H100. B: U40 with 40-μm-diameter. C: U25 with 25-μm-diameter D: 370K with 20-μm-diameter.

#### 2.3 Fluorescence intensity of the U25

Fluorescence intensity of the whole U25 was represented by five regions as shown in Figure S2A. The fluorescence intensities of the five regions from 1-5 were monitored from 0 h-8 h, the CAP analyte was added for the competition analysis at 4 h (Figure S2B). The average gray values of the fluorescence intensities of these five regions from

0 h-8 h was analyzed by the Cellcolecter$^{TM}$ platform shown in Figure S2C.

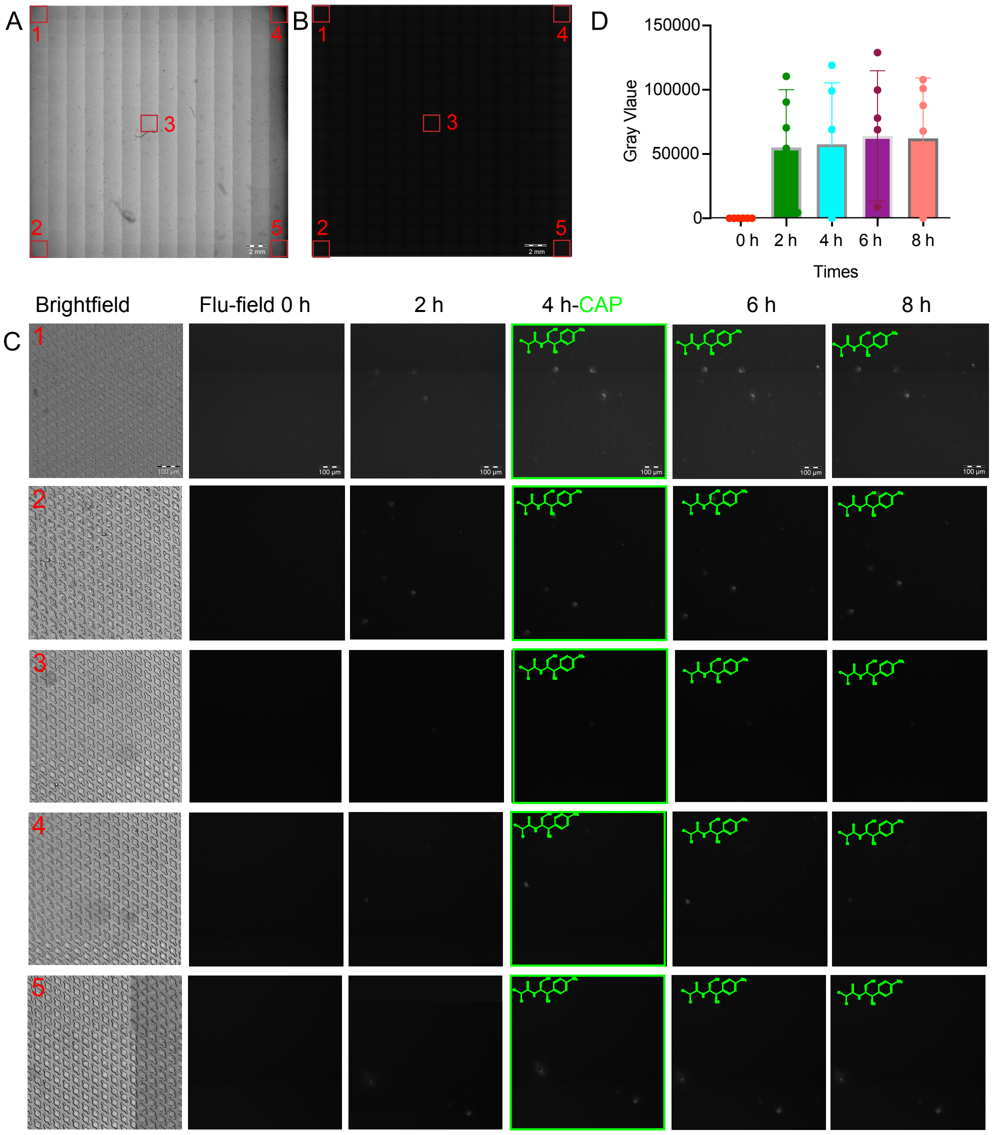


Alt Text: The images fluorescence intensity of five regions represented the whole U25 from 0 h-8 h.

Figure S2. Fluorescence intensity of the whole U25 represented by five regions from 0 h-8 h. A: The brightfield of the whole U25. The red boxes are the chosen five regions. B: The fluorescence field (flu-field) of the while U25. The red boxes are the chosen five regions. C: The detailed fluorescence intensity of the five regions from 0 h-4 h. First column is the brightfield of the region 1-5; second to fifth column are the flu-field from 0 h-8 h, at the time of 4 h, the CAP analytes was added for the competition analysis. D: Histogram analysis of the average gray value of the fluorescence intensity of the 5 regions.

2.4 Preparation of single ACS antibody gene

The CAP-specific single ACS was picked by the tips with 20 μm inner diameter in the microfluidic cell picking robot of the Cellcolecter™ platform. Figure S3 showed the ACS before picking and after picking in the nanowell (red circle).

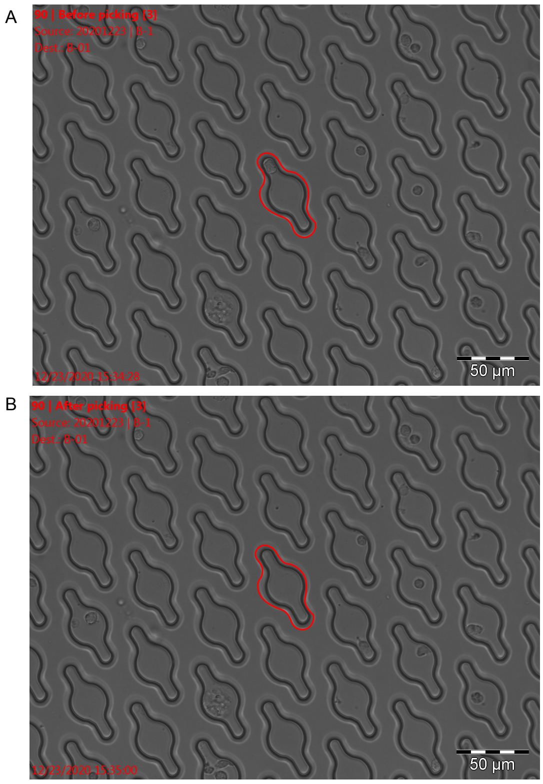


Alt Text: Upper is the image of in U25 nanowell with single ACS before picking. Lower is the image of U25 nanowell with single ACS after picking.

Figure S3. The image of single ACS before picking and after picking. A: Before picking. B: After picking.

Then the picked CAP-specific ACS was lysed, the VH and VL of the the RmAb3 was amplified with the specific primers showed in Table S2. The agarose gel electrophoresis analysis of the VH and VL of 25 retrieved ACSs was shown in Figure S4.

Table S2. Primers of rabbit VH and VL.

| Primers | Sequences |
|---|---|
| RVHF1 | CAGTCGGAGGAGTCCRGG |
| RVHF2 | CAGTCGAAGGAGTCCGAG |
| RVHF3 | CAGTGGAGGAGTCCGGG |
| RVHF4 | CAGSAGTGRTGGAGTCCGG |
| RVHB | TCACCACGCTGCTCAGCGAGT |
| RVKF1 | GAGCTCGTGGACCCAGACTCCA |
| RVKF2 | GAGCTCGAGACCCAGACTCCA |
| RVKB | GGAAGAGGAGGACAGTAGGTGCAACTGGATCCCT |
| RVLF1 | GAGCTCTGACTCAGTCGCCCTC |
| RVLB1 | GCCTGGGTCAGCTGGGTCCC |

R=A/G, Y=C/T, M=A/C, K=G/T, S=C/G, W= A/T, H= A/C/T, B= C/G/T, V=A/C/G, D=A/G /T, N=A/C/G/T.

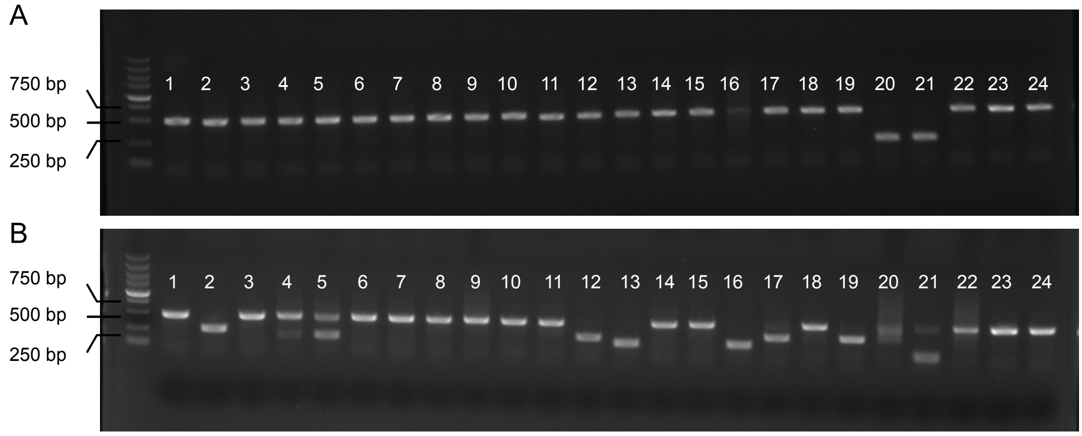


Alt Text: Upper is the image of VH agarose gel electrophoresis analysis. lower is the image of VL agarose gel electrophoresis analysis

Figure S4. The agarose gel electrophoresis analysis of the antibody gene amplified from the retrieved 25 ACSs. A: VL. B: VH.

#### 2.5 Characterization of the antibody from the isolated ACSs

The antibody dilution curves of ELISA based on 16 RmAbs with paired VH and VL were established to analyze the titer of the RmAbs (Figure S5A). The standard curves of icELISA based on the five RmAbs that bond to the CAP-BSA in the ELISA result were established to analyze the $IC_{50}$ to CAP (Figure S5B).

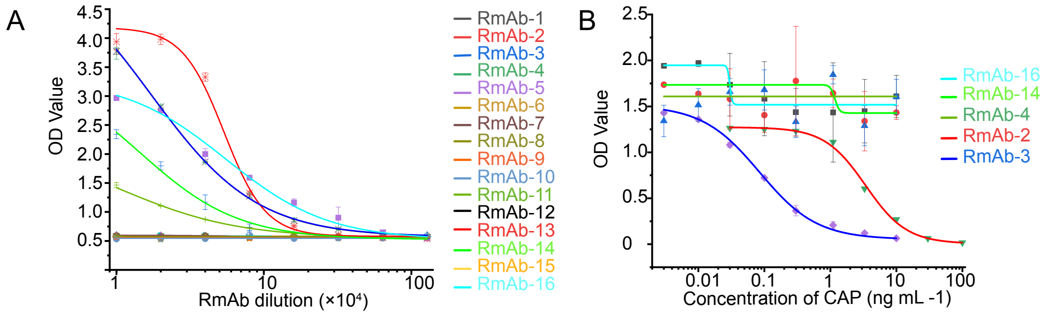


Alt Text: Left is the line chart of the ELISA. The right is the line chart of the icELISA.

Figure S5. Characterization the RmAbs. A: Antibody dilution curves of the ELISA. B: Standard curves of the icELISA for CAP.

The non-denaturing gel electrophoresis showed that the total the RmAb3 is about 154.16 kDa, the denaturing gel electrophoresis showed that the heavy chain of the RmAb3-3 is 54.2 kDa and light chain of the RmAb3-3 is 24.4 kDa (Figure S6).

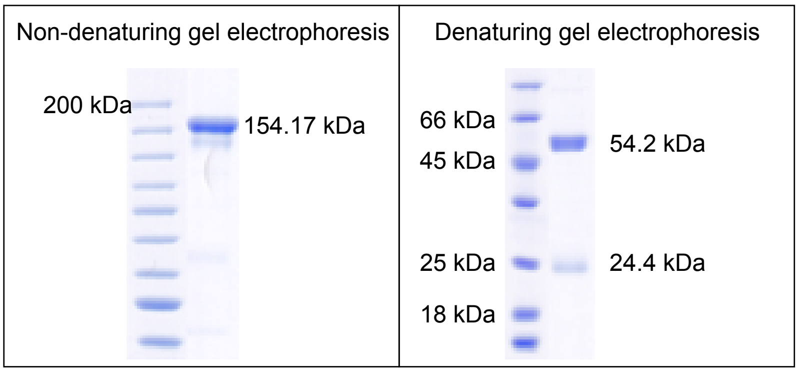


Alt Text: Left is the non-denaturing gel electrophoresis of the RmAb3. The right is denaturing gel electrophoresis the RmAb3.

Figure S6. Polyacrylamide gel electrophoresis analysis of the RmAb3. A: Non-denaturing gel electrophoresis. B: Denaturing gel electrophoresis.

2.6 Characterization of the CAP-specific MmAb

The MmAb produced in our other study (unpublished data) was analyzed as a comparation for the RmAb3. Besides sodium strength, the concentration of methanol and acetonitrile, and pH value of the assay buffer showed a significant influence on CAP and MmAb reactions, which were represented by $IC_{50}/B_0$ (Figure S7A, S7B and S7C). Figure S7D shows the optimized sodium strength of the icELISA based on the MmAb is physiological salt solution of 0.145 M. Tm1 of the MmAb is 74.5°C and Tagg of the MmAb is 74.6°C (Figure S7E). Figure S8F and S9G shows the optimized sodium strength of the icELISA based on the MmAb is physiological salt solution. Figure S7H shows the affinity of MmAb with the $IC_{50}$ of 0.28 ng $mL^{-1}$.

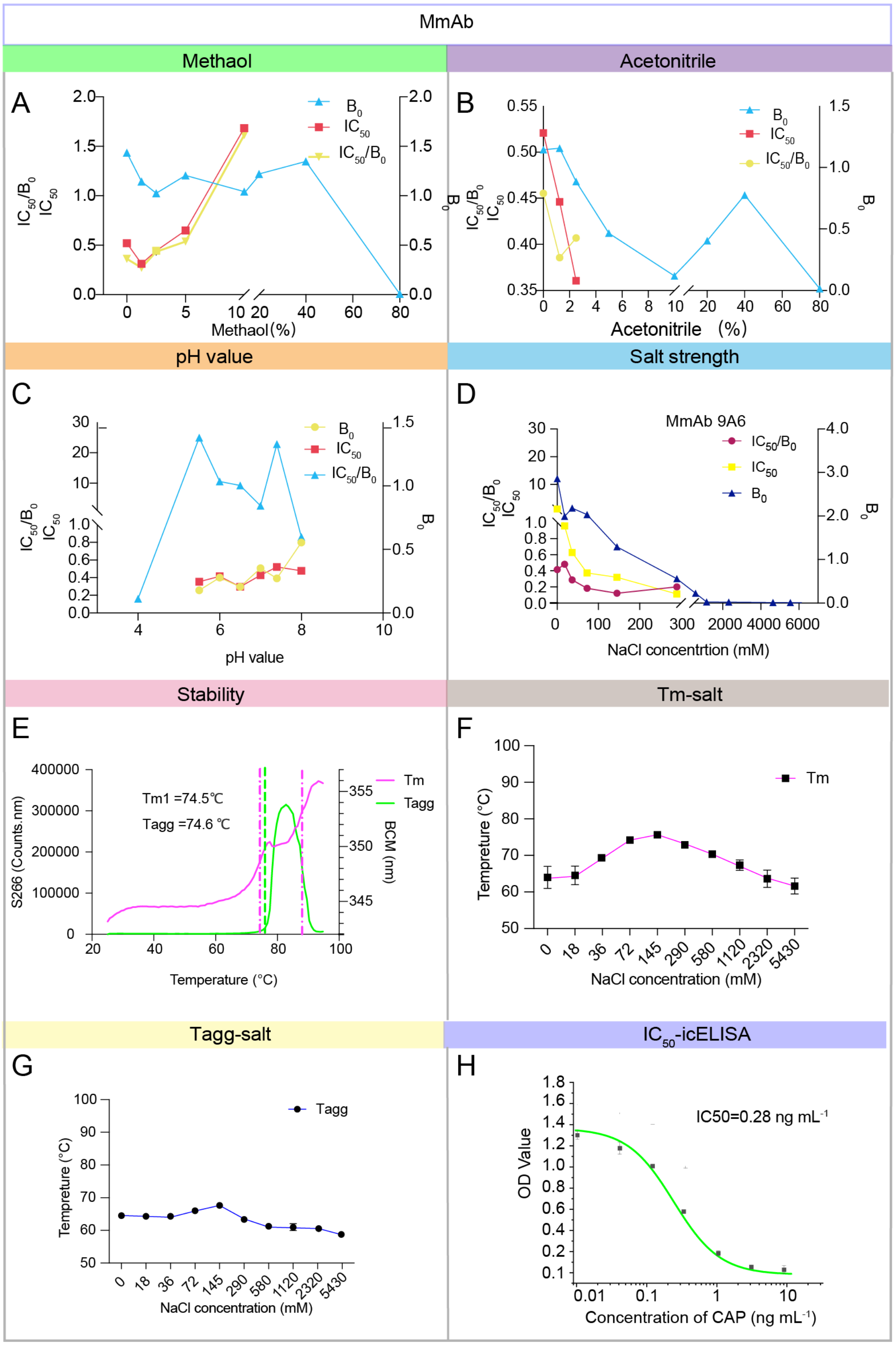


Alt Text: The line charts of the optimization of the methanol, acetonitrile, and pH in the icELISA based on MmAb.

Figure S7. Optimization of the methanol, acetonitrile, and pH in the icELISA based on

MmAb. A: Methanol. B: Acetonitrile. C: pH value. D: The sodium strength. E: Stability analysis of the MmAb. F: The Tm analysis of MmAb at different sodium strength from 0 M -5.43 M. G: The Tagg analysis of MmAb at different sodium strength from 0 M-5.43 M. H: Affinity analysis the MmAb.

2.7 Homology modeling and molecular docking of the RmAb3-Fv-CAP

The homology modeling was based on the sequence showed in Table S3. The type and distribution of residues in the RmAb3 VH and VL are shown in Table S4. In this study, the 3D structures of the RmAb3-Fv were predicted by homology modelling to analyze the binding sites of the antibody. To ensure accurate modeling, five antibody crystal structures with higher similarity (> 88.3%) and greater identity (> 71.1%) were chosen for multiple overlaps according to previous research. Then, CDR regions were optimized based on the IMGT database. Ramachandran plots and Profile-3D were applied to verify predicted torsion angles in the Fv and evaluate the fitness of the protein sequence to ensure rationality. The Ramachandran plot analysis of the constructed model of the RmAb3-Fv showed that 96.7% domain residues were located in the allowed region (Figure S8A), meeting the requirement that a high-quality model with the expected residues with over 90% in the allowed region[1]. Profile-3D analysis indicated that the verification scores of the the RmAb3-Fvs were 110.4 , for slightly higher than the expected high scores of 101.8 [2]. The residue verification score valus shown in Figure S8B is all above 0, indicating that all residues in the model are valid. After the docking simulation, the complexes of Fv residues with the best docking scores of 107.4, was obtained and analyzed.

Table S3. Variable region sequence of the RmAb3.

| mAbs | | Sequence |
|---|---|---|
| RmAb3 | VH [a] | QSVEESGGRLVTPGTPLTLTCTASGFSSLNYYMTAPQQAPGKGLEWIGAINYTTITYYASWAKGRFTISKTSTTVDLRITSPTTEDTFAYTCARGAGSSDDTMGYYFNIWGPGTLVTVSS |
| | VL [b] | ELVMTQTPASVSAAVGGTVTINCQASDNYSNIIYILAWYKPQQGQRPRLLIFGASTLESGVPSRFKGSGSGTEFTLTISCAADTYYCQCTDYRGSSDNVFGGGTEVVVK |

[a] VH is the variable region sequence of heavy chain. [b] VL is the variable region sequence of light chain.

Table S4. Amino acid analysis of the VH and VL of the RmAb3 and MmAb.

| Amino acid | RmAb3 VH | | RmAb3 VL | | MmAb VH | | MmAb VL | |
|---|---|---|---|---|---|---|---|---|
| | NO. | Percentage (%) | NO. | Percentage (%) | NO. | Percentage (%) | NO. | Percentage (%) |
| Ala (A) | 9 | 7.5% | 8 | 7.3% | 5 | 4.2% | 5 | 4.5% |
| Arg (R) | 4 | 3.3% | 4 | 3.7% | 4 | 3.4% | 3 | 2.7% |
| Asn(N) | 3 | 2.5% | 4 | 3.7% | 8 | 6.8% | 3 | 2.7% |
| Asp(D) | 4 | 3.3% | 5 | 4.5% | 4 | 3.4% | 4 | 3.6% |
| Cys (C) | 2 | 1.7% | 4 | 3.7% | 2 | 1.7% | 2 | 1.8% |
| Gln (Q) | 3 | 2.5% | 6 | 5.5% | 7 | 5.9% | 5 | 4.5% |
| Glu (E) | 4 | 3.3% | 5 | 4.5% | 3 | 2.5% | 5 | 4.5% |
| Gly (G) | 13 | 11.0% | 12 | 11.0% | 11 | 9.3% | 9 | 8.1% |
| His (H) | 0 | 0.0% | 0 | 0.0% | 2 | 1.7% | 1 | 0.9% |
| Ile (I) | 6 | 5.0% | 6 | 5.5% | 7 | 5.9% | 3 | 2.7% |
| Leu (L) | 7 | 5.8% | 6 | 5.5% | 10 | 8.5% | 12 | 10.8% |
| Lys (K) | 3 | 2.5% | 3 | 2.8% | 5 | 4.2% | 7 | 6.3% |
| Met(M) | 2 | 1.7% | 1 | 0.9% | 0 | 0% | 2 | 1.8% |
| Phe (F) | 4 | 3.3% | 4 | 3.7% | 5 | 4.2% | 3 | 2.7% |
| Pro (P) | 6 | 5.0% | 4 | 3.7% | 4 | 3.4% | 5 | 4.5% |
| Ser (S) | 13 | 10.8% | 12 | 11.0% | 13 | 11.0% | 18 | 16.2% |
| Thr (T) | 21 | 17.5% | 11 | 10.1% | 11 | 9.3% | 8 | 7.2% |
| Trp(W) | 3 | 2.5% | 1 | 0.9% | 4 | 3.4% | 2 | 1.8% |
| Tyr (Y) | 8 | 6.7% | 6 | 5.5% | 8 | 6.8% | 7 | 6.3% |
| Val (V) | 5 | 4.2% | 9 | 8.3% | 5 | 4.2% | 7 | 6.3% |

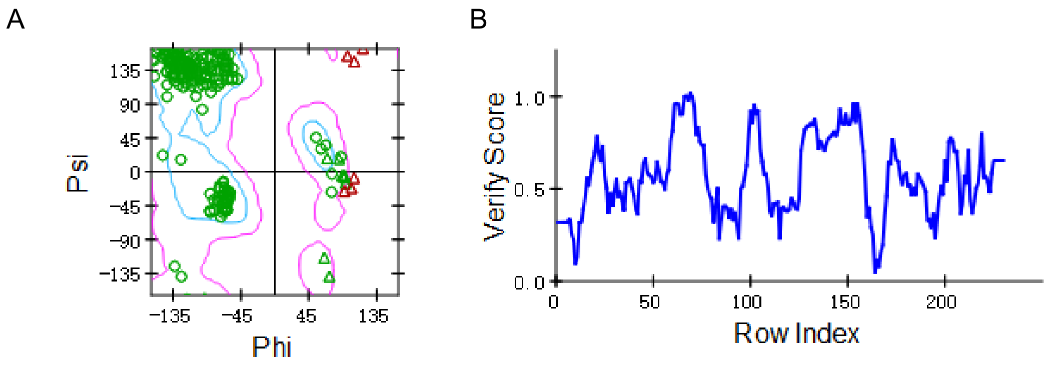


Alt Text: The verification of the RmAb3-Fv homology model.

Figure S8. The Ramachandran plot and Profile-3D analysis of the RmAb3-Fv homology model. A: Ramachandran plot analysis of the RmAb3-Fv homology model. B: Profile-3D analysis of the RmAb3-Fv homology model.

2.9 MD simulation of the RmAb3-Fv-CAP and the MmAb-Fv-CAP

The structures of the RmAb3-Fv-CAP and the MmAb-Fv-CAP complexes before and after MD at different concentration of NaCl system of control salt solution, physiological salt solution and saturated salt solution are shown in Figure S9. Root mean square deviation (RMSD) was analyzed to evaluated the stability of the whole structure and the constructed system of the mAb Fv-CAP complex (Figure S10A and Figure S11A). Solvent accessible surface area (SASA) analysis was conducted to evaluate the surface structure changes of the RmAb3-Fv-CAP complex and the MmAb-Fv-CAP complex (Figure S10B and Figure S11B). Radius of gyration (Rg) was used to assess the stability of the whole structure (Figure S10C and Figure S11C), root mean square fluctuation (RMSF) analyses are used to evaluate the flexible of the RmAb3-CAP complex and the MmAb-Fv-CAP complex (Figure S10D and Figure S11D).

Finally, inter-hydrogen bonds are analyzed to judge the interaction between the RmAb3-Fv/ MmAb-Fv and CAP (Figure S10E and Figure S11E).

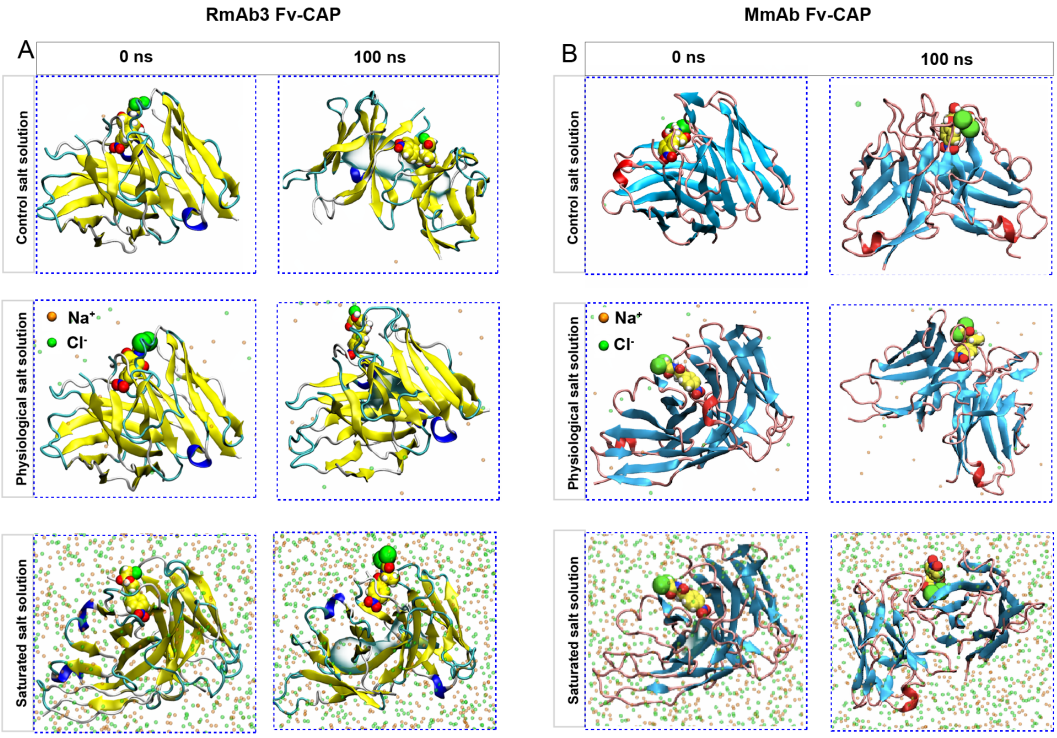


Alt Text: The MD analysis of RmAb3-Fv-CAP and the MmAb-Fv-CAP complexes at different concentration of NaCl system.

Figure S9. The structures of the RmAb3-Fv-CAP and the MmAb-Fv-CAP complexes at different concentration of NaCl system before and after MD. A: The structures of the RmAb3-Fv-CAP complex at control salt solution, physiological salt solution and saturated salt solution before and after MD. Left column is before MD at 0 s, right column is after MD at 100 s. B: The structures of the MmAb-Fv-CAP complex at control salt solution, physiological salt solution and saturated salt solution before and after MD. Left column is before MD at 0 s, right column is after MD at 100 s.

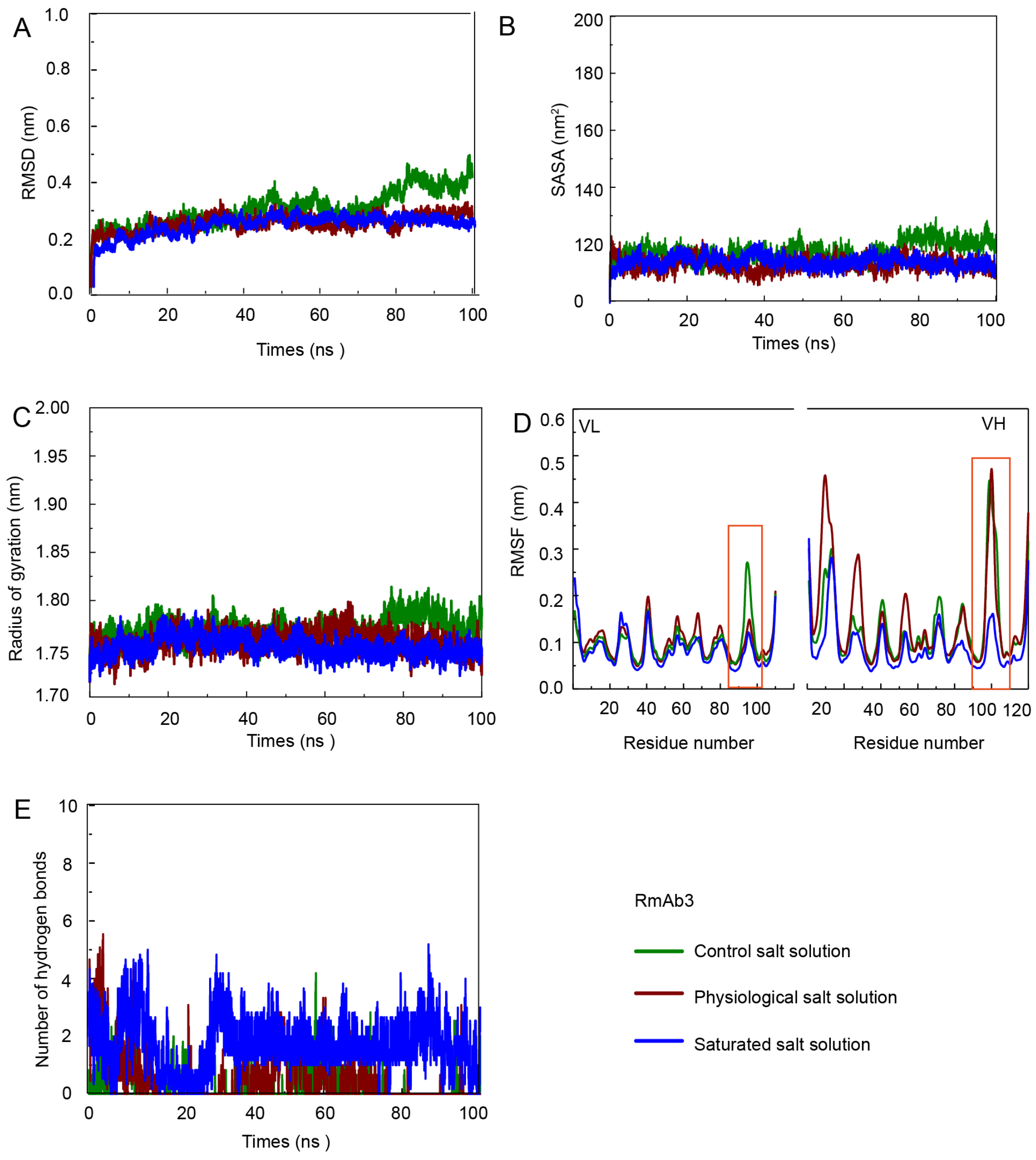


Alt Text: The post stability analysis of RmAb3-Fv-CAP complexe at different concentration of NaCl system.

Figure S10. Characterizations of the MD of the RmAb3-Fv-CAP complex. A: RMSD analysis of the the RmAb3-Fv-CAP complex. B: SASA analysis of the the RmAb3-Fv-CAP complex. C: Rg analysis of the the RmAb3-Fv-CAP complex. D: RMSF analysis of the the RmAb3-Fv-CAP complex. E: Hydrogen bond analysis of the the RmAb3-Fv-CAP complex.

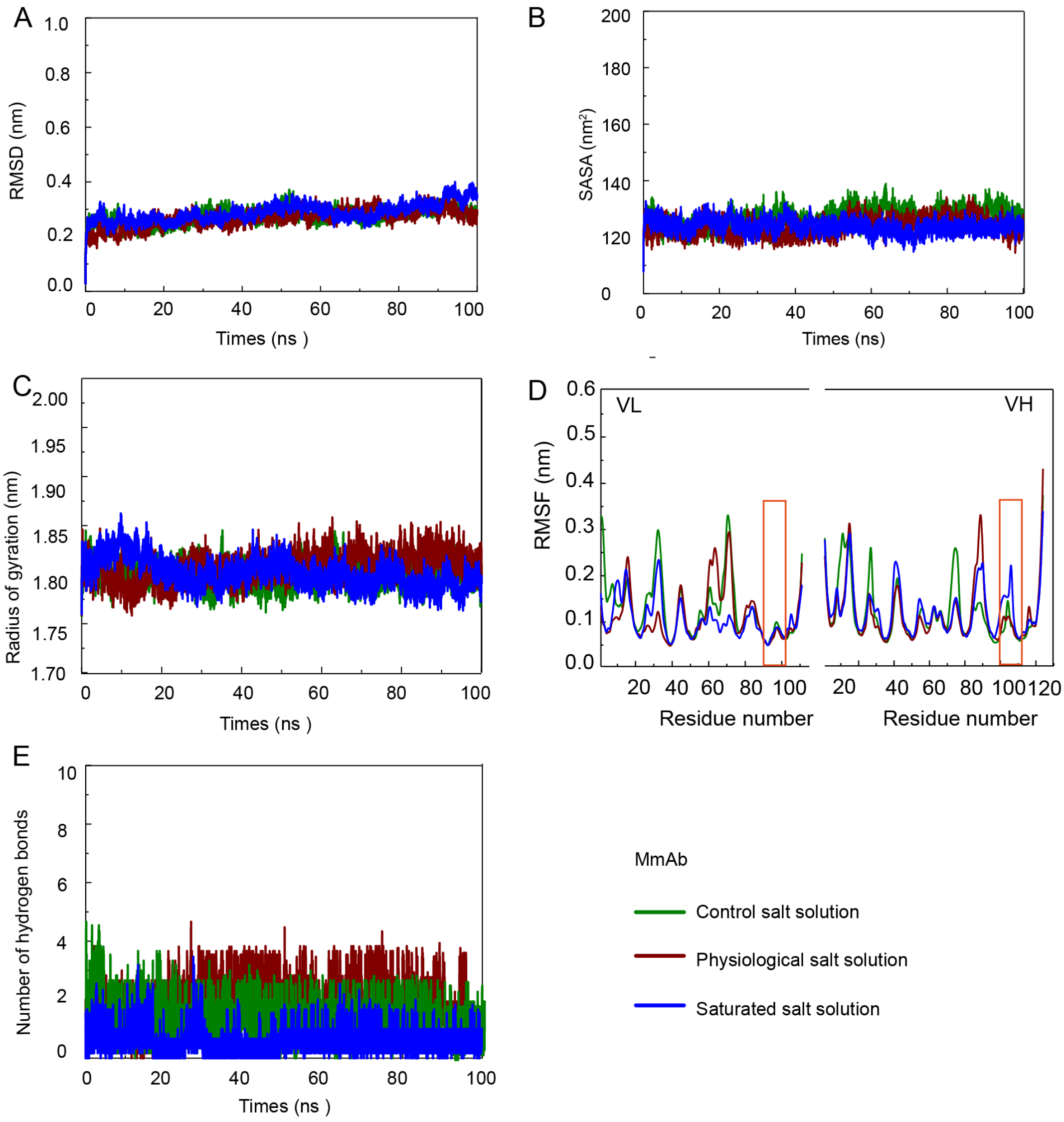


Alt Text: The post stability analysis of the MmAb-Fv-CAP complexe at different concentration of NaCl system.

Figure S11. Characterizations of the MD of the MmAb-Fv-CAP complex. A: RMSD analysis of the the MmAb-Fv-CAP complex. B: SASA analysis of the the MmAb-Fv-CAP complex. C: Rg analysis of the the MmAb-Fv-CAP complex. D: RMSF analysis of the the MmAb-Fv-CAP complex. E: Hydrogen bond analysis of the the MmAb-Fv-CAP complex.

2.11 Immunoassay development and sample evaluation

The development of the icELISA of the RmAb3 was based on the optimum assay buffer. The specify of the the RmAb3 with the lowest $IC_{50}$ was analyzed shown in Table S4. The HPLC-MS/MS results are shown in Table S5.

Table S4. The specificity of the RmAb3.

| Compound | Structure | $IC_{50}$ (ng $mL^{-1}$) | CR [e] |
|---|---|---|---|
| CAP [a] | 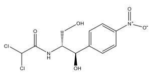 | 0.08 | 100% |
| TAP [b] | 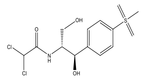 | >1000 | <0.01% |
| FF [c] | 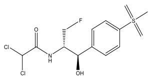 | >1000 | <0.01% |
| FFA [d] | 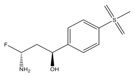 | >1000 | <0.01% |

[a] CAP is the chloramphenicol. [b] TAP is the thiamphenicol. [c] FF is the florfenicol. [d] FFA is the florfenicol amide. [e] CR is cross-reactivity and calculated according to Eq.3.

Table S5. Detection of CAP in positive chicken samples using icELISA and HPLC-MS/MS (N=6).

| Samples | icELISA<br>CAP (μg $kg^{-1}$) | UPLC-MS/MS[a]<br>CAP (μg $kg^{-1}$) |
|---|---|---|
| Chicken 1 | -[b] | 1.262 ± 0.020 |
| Chicken 2 | 0.297 ± 0.000 | 0.349 ± 0.007 |
| Chicken 3 | 0.512 ± 0.014 | 0.602 ± 0.004 |
| Chicken 4 | 1.032 ± 0.007 | 1.180 ± 0.017 |
| Chicken 5 | -[b] | 1.417 ± 0.009 |
| Chicken 6 | - [b] | 5.83 ± 0.009 |
| Chicken 7 | - [b] | 3.005 ± 0.008 |
| Chicken 8 | 0.917 ± 0.003 | 1.004 ± 0.000 |
| Chicken 9 | 0.544 ± 0.000 | 0.532 ± 0.004 |
| Chicken 10 | 0.163 ± 0.002 | 0.154 ± 0.004 |

| | | |
|---|---|---|
| Chicken 11 | 0.152 ± 0.000 | 0.147 ± 0.000 |
| Chicken 12 | 0.618 ± 0.030 | 0.593 ± 0.003 |

[a] The detection method based on the GB/T 22338-2008.
[b] Out of detection range
